# Discovery of Clinically Relevant Fusions in Pediatric Cancer

**DOI:** 10.1101/2021.03.11.435013

**Authors:** Stephanie LaHaye, James R. Fitch, Kyle J. Voytovich, Adam C. Herman, Benjamin J. Kelly, Grant E. Lammi, Saranga Wijeratne, Samuel J. Franklin, Kathleen M. Schieffer, Natalie Bir, Sean D. McGrath, Anthony R. Miller, Amy Wetzel, Katherine E. Miller, Tracy A. Bedrosian, Kristen Leraas, Kristy Lee, Ajay Gupta, Bhuvana Setty, Daniel R. Boué, Jeffrey R. Leonard, Jonathan L. Finlay, Mohamed S. Abdelbaki, Diana S. Osorio, Selene C. Koo, Daniel C. Koboldt, Vincent Magrini, Catherine E. Cottrell, Elaine R. Mardis, Richard K. Wilson, Peter White

**Affiliations:** The Steve and Cindy Rasmussen Institute for Genomic Medicine, Nationwide Children’s Hospital, Columbus, OH; Division of Hematology, Oncology, Blood and Marrow Transplant, Nationwide Children’s Hospital, Columbus, OH; Department of Pediatrics, The Ohio State University, Columbus, OH; Department of Pathology, The Ohio State University, Columbus, OH; Department of Pathology, Nationwide Children’s Hospital, Columbus, OH; Section of Neurosurgery, Nationwide Children’s Hospital Columbus, OH

**Keywords:** transcriptomics, genomics, pediatric neoplasms, gene fusions, cancer, RNA-Seq

## Abstract

**Background:** Pediatric cancers typically have a distinct genomic landscape when compared to adult cancers and frequently carry somatic gene fusion events that alter gene expression and drive tumorigenesis. Sensitive and specific detection of gene fusions through the analysis of next-generation-based RNA sequencing (RNA-Seq) data is computationally challenging and may be confounded by low tumor cellularity or underlying genomic complexity. Furthermore, numerous computational tools are available to identify fusions from supporting RNA-Seq reads, yet each algorithm demonstrates unique variability in sensitivity and precision, and no clearly superior approach currently exists. To overcome these challenges, we have developed an ensemble fusion calling approach to increase the accuracy of identifying fusions.

**Results:** Our ensemble fusion detection approach utilizes seven fusion calling algorithms: Arriba, CICERO, FusionMap, FusionCatcher, JAFFA, MapSplice, and STAR-Fusion, which are packaged as a fully automated pipeline using Docker and AWS serverless technology. This method uses paired end RNA-Seq sequence reads as input, and the output from each algorithm is examined to identify fusions detected by a consensus of at least three algorithms. These consensus fusion results are filtered by comparison to an internal database to remove likely artifactual fusions occurring at high frequencies in our internal cohort, while a “known fusion list” prevents failure to report known pathogenic events. We have employed the ensemble fusion-calling pipeline on RNA-Seq data from 229 patients with pediatric cancer or blood disorders studied under an IRB-approved protocol. The samples consist of 138 central nervous system tumors, 73 solid tumors, and 18 hematologic malignancies or disorders. The combination of an ensemble fusion-calling pipeline and a knowledge-based filtering strategy identified 67 clinically relevant fusions among our cohort (diagnostic yield of 29.3%), including *RBPMS-MET, BCAN-NTRK1*, and *TRIM22-BRAF* fusions. Following clinical confirmation and reporting in the patient’s medical record, both known and novel fusions provided medically meaningful information.

**Conclusions:** Our ensemble fusion detection pipeline offers a streamlined approach to discover fusions in cancer, at higher levels of sensitivity and accuracy than single algorithm methods. Furthermore, this method accurately identifies driver fusions in pediatric cancer, providing clinical impact by contributing evidence to diagnosis and, when appropriate, indicating targeted therapies.

## Background

Globally, there are approximately 300,000 pediatric and adolescent cases of cancer diagnosed each year [1, 2]. While advances in medicine have led to a drastic improvement in 5-year overall survival rates (up to 84% in children under 15), pediatric cancer remains the most common cause of death by disease in developed countries [3, 4]. Pediatric cancers are defined by a distinct genomic landscape when compared to adult cancers, which includes an overall low number of somatic single nucleotide variants, common driver fusions and epigenetic changes that drive a specific transcriptional program. Pediatric cancers are often considered embryonic in origin and demonstrate a significant germline predisposition component approaching 10% [5–7].

Many pediatric tumors contain gene fusions resulting from the juxtaposition of two genes (**Additional File 1: Figure S1**) [6]. Fusions typically occur through chromosomal rearrangements, and often lead to dysregulated gene expression of one or both gene partners [8–11]. Fusions can also generate chimeric oncoproteins, wherein functional domains from both genes are retained, often leading to aberrant and strong activation of nonspecific downstream targets [12]. The alterations in gene expression and activation of downstream targets induced by fusions are considered to be oncogenic events in pediatric cancer and increasingly may indicate response to specific targeted therapies.

The identification of an oncogenic fusion can provide medically meaningful information in the context of diagnosis, prognosis, and treatment regimens in pediatric cancers. Fusions may provide diagnostic evidence for a specific histological subgroup. For example, *EWSR1-FLI1* fusions are highly associated with Ewing sarcoma, while the presence of a *C11orf95-RELA* fusion aids in subgrouping supratentorial ependymomas [12]. The detection of certain fusions, such as *BCR-ABL* in acute lymphocytic leukemia, can be used as a surrogate for residual tumor load and treatment response [13]. Fusions may also provide prognostic indication, such as *KIAA1549-BRAF* in low grade astrocytomas, which have a more favorable outcome compared to non-*BRAF* fused tumors [14,15]. In addition, fusions that involve kinases can present therapeutic targets, including *FGFR1-TACC1, FGFR3-TACC3, NPM1-ALK*, and *NTRK* fusions [2,12,16-19].

However, regardless of the clear clinical benefits of characterizing fusion events in a given patient’s tumor, accurate identification of fusions from next generation sequencing DNA data alone is not straightforward and they often go undiscovered. In particular, many fusions are not detectable by exome sequencing (ES) due to breakpoint locations that frequently occur in non-coding or intronic regions which may not have corresponding capture probes. Even whole genome sequencing (WGS) NGS data has proved difficult to evaluate complex rearrangements resulting in gene fusions due to a high false positive rate and due to the limitations of short read lengths [20, 21]. By contrast, next-generation RNA sequencing data, or RNA-Sequencing (RNA-Seq), offers an unbiased data type suitable for fusion detection, while also providing information about the expression of fusion transcripts, including multiple isoforms, and fusions that occur due to aberrant splicing events [22, 23],

While RNA-Seq is a powerful tool for fusion detection, it is not without its limitations. Notably, there is currently a major deficit in our ability to accurately identify fusions in spite of having many computational approaches available. Here, consistently identifying gene fusion events with high sensitivity and precision using one algorithm is unlikely and this is of critical importance in a clinical diagnostic setting [12]. Computational approaches that have been tuned for high sensitivity are limited by also calling numerous false positives, requiring extensive manual review of data, while those with a low false discovery rate (FDR) often miss true positives due to over-filtering [12]. To overcome these complications of sensitivity and specificity, we have employed an ensemble pipeline, which merges results from seven algorithmic approaches to identify, filter and output prioritized fusion predictions.

Another common issue encountered in fusion prediction is the identification of likely non-pathogenic fusions, due both to read-through events and fusions occurring in non-disease involved (normal) genomes. [12, 24, 25] We addressed these sources of false positivity through the implementation of a filtering strategy that removes known normal fusions and RNA transcription read-through events, based on internal frequency of detection and location of chromosomal breakpoints. Lastly, to prevent over-filtering and inadvertent removal of previously described known pathogenic fusion events, we have developed and continually update a list containing known pathogenic fusion partners, that will return any data-supported fusions to the output list of prioritized fusion results for further evaluation.

The ensemble fusion detection pipeline outperformed all single algorithm methods we evaluated, achieving high levels of sensitivity, while simultaneously minimizing false positive calls and non-clinically relevant fusion predictions. Here, we describe our ensemble fusion detection approach, its performance on commercial control reference standards with known fusions, and its implementation on a pediatric cohort consisting of rare, treatment refractory, or relapsed cancers and hematologic diseases. Utilization of our ensemble approach resulted in a diagnostic yield of approximately 30% in our cohort, identified novel fusion partners, and has provided diagnostic information and/or targeted treatment options for this patient population.

## Results

### Development and optimization of ensemble pipeline on a control reference standard

Identification of gene fusions through the use of a single algorithm is often associated with low specificity and poor precision [12]. Given prior literature supporting multi-algorithmic approaches to improve upon these deficits, we studied the intricacies of several fusion detection algorithms, and applied a defined set of algorithms with desired properties, aimed at detecting true positive fusions while minimizing false positive fusions [25–27]. After evaluating each algorithm’s output, we developed our ensemble fusion detection pipeline that combines output consensus calls from seven different computational approaches (**Figure 1A**), calculates the concordant fusion partners and breakpoints, and filters this output list based on internal frequency, reads of evidence, and breakpoint location. A list of known pathogenic fusions rescues any known pathogenic fusion gene partners with suitable algorithmic and read support for further evaluation (**Additional File 1: Table S3**).

**Figure 1.**
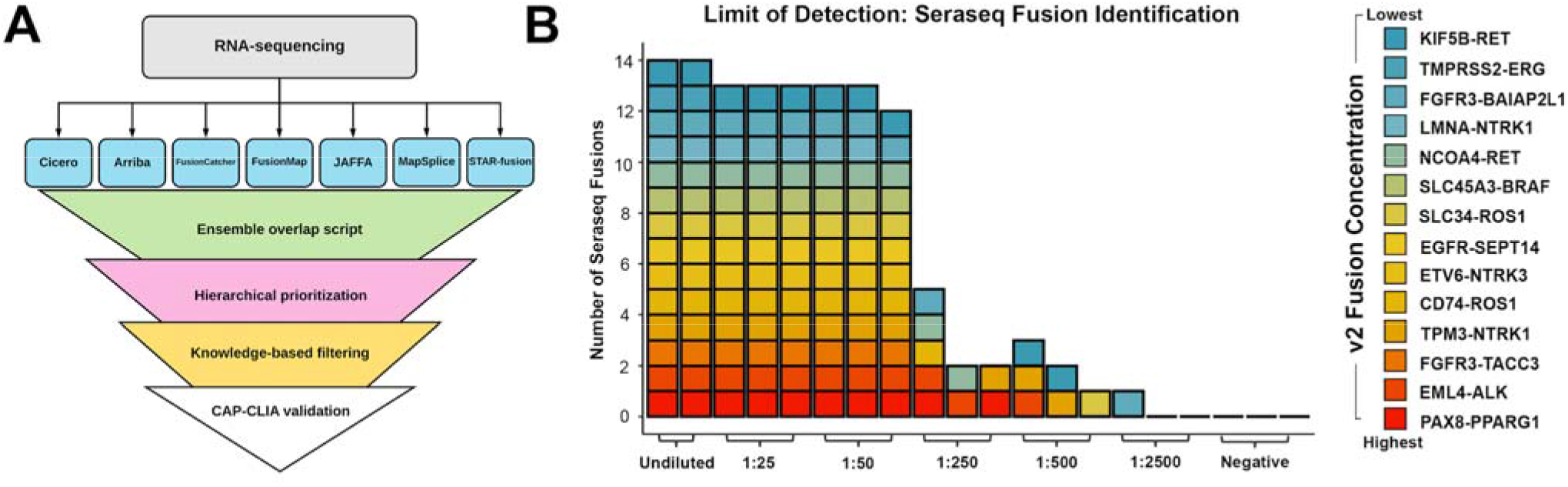
The ensemble fusion detection pipeline identifies true positive fusions. **A)** The ensemble approach identifies fusions in RNA-Seq data by overlapping results from Arriba, CICERO, FusionCatcher, FusionMap, JAFFA, MapSplice, and STAR-Fusion. It hierarchically prioritizes and filters the fusions utilizing an in-house PostgreSQL database and knowledge base, prior to producing an output list of predicted fusions. In many cases, detected fusions were orthogonally tested by clinical confirmation in order to return a medically meaningful result. **B)** The ensemble pipeline was tested on a dilution series of a reference control reagent (SeraCare) to determine sensitivity and limit of detection. We optimized the pipeline using the undiluted reference control reagent, identifying that by requiring ≥3 callers to have overlap for a detected fusion, and by utilizing filtering of known false positive fusion calls and cross-referencing a list of known fusions, all 14 fusions were identified. Colors representing different fusions present in the SeraSeq v2 reagent are ordered by their absolute proportions. We then applied the optimized pipeline to the dilution series, showing that the numbers of identified fusions were reduced in serial dilutions, and no fusions were identified in the negative control.

To optimize the approach, we utilized a reference standard from a commercial provider (Seraseq Fusion RNA, SeraCare, Milford, MA), containing synthetic RNAs representing 14 cancer-associated fusions in varying proportions (**Additional File 1: Tables S1 and S2**). Data generated from these RNA-Seq libraries, performed as replicates for a range of dilutions, were analyzed using the ensemble pipeline. We compared the output derived from a consensus of two or more callers to that from a consensus of three or more callers by calculating sensitivity (# of Seraseq fusions identified)/(14 possible Seraseq fusions), and precision (# of Seraseq fusions identified)/(# of total fusions identified) prior to filtering or known fusion list comparison. The undiluted reference standard with consensus of at least two callers, had a sensitivity of 100% and precision of 36.36%. Inclusion of the knowledgebase filtering step reduced the sensitivity to 85.71% while increasing the precision to 77.42%, and the known fusion list rescue step increased sensitivity to 100% and precision to 80% (**Additional File 1: Table S5, Figure S6A**). By increasing the consensus requirement to three callers, rather than just two, the prefiltered sensitivity was 100% and precision was 93.33%. Inclusion of the filtering step reduced the sensitivity to 85.71% while increasing the precision to 100%, and known fusion list rescue increased sensitivity to 100% and precision to 100% (**Table 2; Additional File 1: Figure S6A**). The inclusion of the known fusion list prevented the removal of known Seraseq fusions, due to too few reads of evidence or number of callers providing support, as well as a single Seraseq fusion, *EML4-ALK,* which was present at an artificially high frequency in our database (24.7%) due to false positive calls by FusionCatcher. Implementation of the known fusion list led to sensitivity scores of 100% for both levels of caller consensus. The individual fusion detection algorithms ranged in sensitivity and precision, and while certain algorithms are able to maintain high levels of sensitivity in addition to moderate levels of precision, such as STAR-Fusion (sensitivity = 100%, precision = 43.75%), others such as FusionCatcher (sensitivity = 92.86%, precision = 4.34%) and CICERO (sensitivity = 100%, precision 1.06%) had high levels of sensitivity with very low precision levels (**Table 2; Additional File 2: Table S5**). When considering the overall results from undiluted and serial dilutions of the reference standard, the required overlap of at least three callers, with filtering and utilization of the known fusion list, led to significantly fewer total fusions identified compared to two consensus callers (p = 1.86E-07)(**Table 2; Additional File 1: Figure S6B, Table S6**). The ensemble pipeline results obtained from various reference standard dilutions, with a minimum of three callers in consensus, using filtering and known fusion list rescue are shown (**Figure 1B; Additional File 2: Table S5**). The optimized ensemble pipeline, consisting of a consensus of three callers, filtering, and the known fusion list, maintained high levels of sensitivity, (at least 90.48%), while maintaining 100% precision as low as the 1:50 dilution of the reference standard (**Additional File 2: Table S5**). In addition to the high levels of sensitivity and precision, the total number of fusions identified by this optimized ensemble pipeline in undiluted and diluted samples was significantly fewer than the number identified by individual fusion detection algorithms, including STAR-Fusion (p = 1.77E-12), CICERO (p = 3.39E-14) and FusionCatcher (p = 1.00E-08) (**Additional File 1, Table S6**). These results highlights the removal of false positive fusions, which includes artifactual and benign fusion events, and subsequent reduction in manual evaluation requirements (**Additional File 1: Figure S6C,D**). Notably, we only considered the 14 Seraseq synthetic fusions as true positives. While fusions may exist within the GM24385 cell line, in the optimized ensemble approach all of these fusions were filtered out due to either high frequency across our cohort or supporting read evidence below our minimum threshold, suggesting that they are likely to be artifactual in nature.

**Table 2.** Improved precision in fusion detection, utilizing Seraseq controls, achieved through utilization of the ensemble pipeline. Data shown is from undiluted Seraseq v3 RNA-Seq, experiments performed in duplicate, averages are shown. Individual algorithms are listed by precision, in descending order. Seraseq fusions identified (true positive) are out of a possible 14 fusions.

| Algorithm | Total Fusions Identified | Seraseq Fusions Identified | Sensitivity | Precision |
| --- | --- | --- | --- | --- |
| Arriba | 23.5 | 13 | 92.9% | 55.3% |
| MapSplice | 22 | 12 | 85.7% | 54.6% |
| STAR-fusion | 32 | 14 | 100.0% | 43.6% |
| FusionMap | 30 | 12.5 | 89.3% | 41.7% |
| FusionCatcher | 299.5 | 13 | 92.9% | 4.3% |
| JAFFA | 470.5 | 12.5 | 89.3% | 2.7% |
| CICERO | 1323 | 14 | 100.0% | 1.1% |
| Ensemble 2 callers | 38.5 | 14 | 100.0% | 36.4% |
| Ensemble 2 callers + filter | 15.5 | 12 | 85.7% | 77.4% |
| Ensemble 2 callers + filter + known fusion list | 17.5 | 14 | 100.0% | 80.0% |
| Ensemble 3 callers | 15 | 14 | 100.0% | 93.3% |
| Ensemble 3 callers + filter | 12 | 12 | 85.7% | 100.0% |
| Ensemble 3 callers + filter + known fusion list | 14 | 14 | 100.0% | 100.0% |

### Implementation of the ensemble approach on an in-house pediatric cancer and hematologic disease cohort

Having demonstrated the efficacy of the optimized ensemble fusion detection pipeline using synthetic fusion samples, we further evaluated the utility of the pipeline on RNA-Seq data obtained from 229 patient samples, obtained from three prospective pediatric cancer and hematologic disease studies at Nationwide Children’s Hospital (NCH) (**Additional File 1: Figure S2**). Our ensemble pipeline identified significantly fewer total predicted fusions post-filtering, compared to all other single callers (**Figure 2A; Additional File 1: Table S7). Applying the known fusion list rescue altered the average number of fusions identified overall, as an average of 3.88 fusions per case were identified by 3 or more callers, while an average of 3.93 fusions were identified by 3 or more callers after applying the known fusion list; a total of 11 fusions were rescued by this approach, of which 1 (*KIAA1549-BRAF*; Additional File 3: Table S8**) was clinically relevant. The retained *KIAA1549-BRAF* fusion was identified by three callers, but was initially filtered out due to too few reads of evidence, possibly due to either low expression, low tumor cellularity or clonality (**Figure 2D**). In total, 67 clinically relevant fusions, identified in 67 different cases, (33 CNS, 7 heme, and 27 solid tumor; **Additional File 1: Figure S7**) were discovered using the optimized ensemble pipeline with automated filtering, including the known fusion list feature, and a consensus of three callers (29.3% of tumors contained a clinically relevant fusion). Regardless of source material, there was a roughly a 30% yield; with clinically relevant fusion identification in 44 of 148 frozen samples (30% yield), 19 of 68 FFPE samples (28% yield), and 4 of 13 other samples (31% yield), which included blood, cerebral spinal fluid, or bone marrow (**Additional File 1: Figure S7**). No single fusion detection algorithm was able to identify all 67 fusions. While JAFFA was the most sensitive algorithm, identifying the most clinically relevant fusions (64 out of 67), it also had one of the highest average numbers of fusions identified per sample, 1409 fusions, indicating a large number of likely false positives (**Figure 2B; Additional File 1: Table S7**). Identified fusions were broken down into 4 types: Interchromosomal Chimeric (n= 30), Intrachromosomal Chimeric (n= 29), Loss of Function (n= 3), and Promoter Swapping (n= 5)(**Figure 2C**). Of the 67 clinically relevant fusions, seven were considered novel events, defined as a gene fusion involving two partners not previously described in the literature at the time of identification (**Figure 2D**). Of the 67 fusions detected, 40 (60%) were identified by all seven callers, 55 (82%) were identified by ≥6 callers, 60 (90%) were identified by ≥5 callers, 64 (96%) were identified by ≥4 callers, and 67 (100%) were identified by ≥3 callers. (**Figure 2E**). One sample experienced an unresolvable failure of FusionMap, likely due to high sequencing read number. Results from the remaining callers, which successfully completed for this sample, were still included in our analysis. These results highlight the ability of the optimized ensemble approach to identify gene fusions with a high level of confidence and a reduced number of false positive predictions, while preventing over-filtering by comparison to a list of known pathogenic fusions.

**Figure 2.**
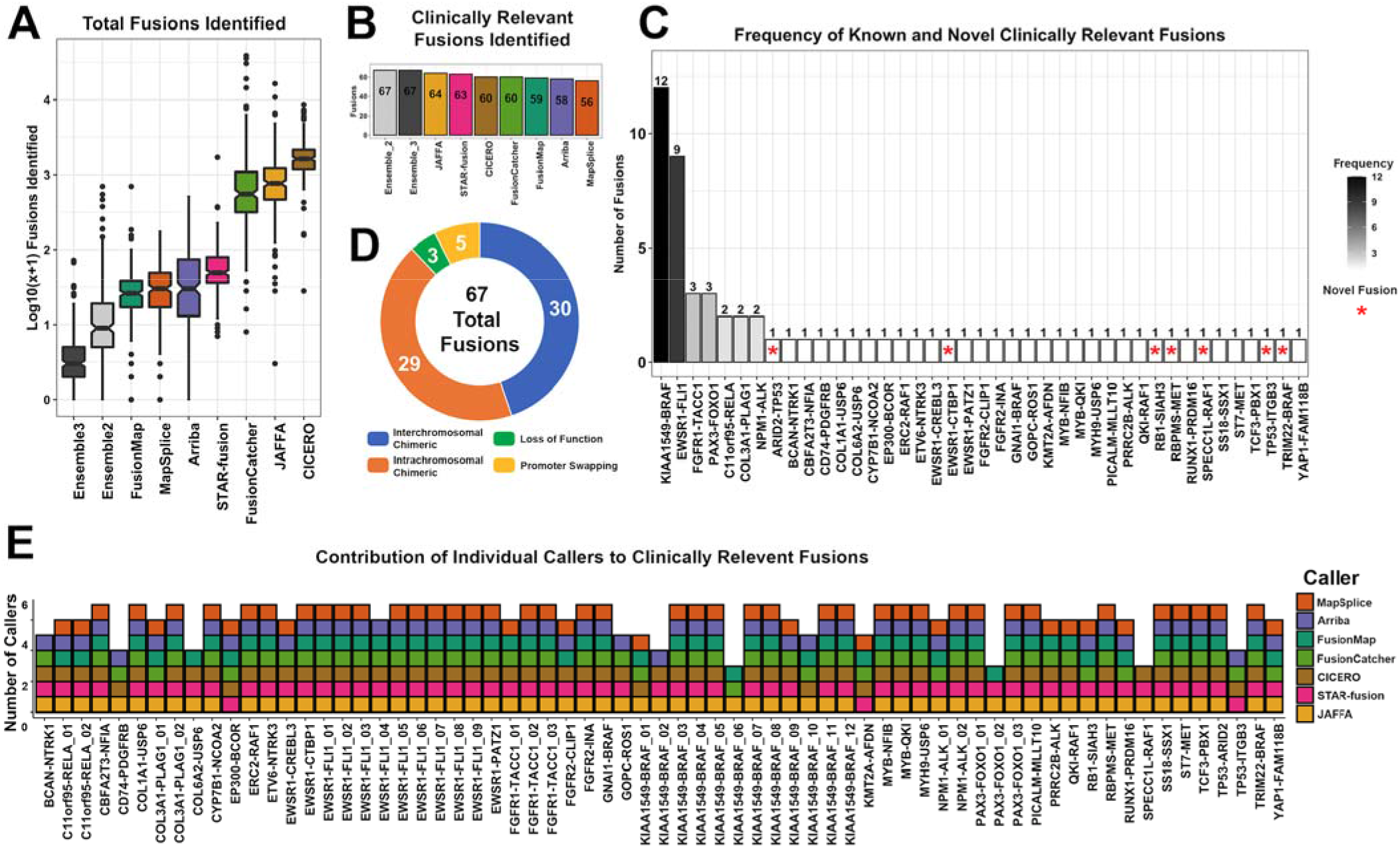
Clinically relevant fusions identified by the ensemble approach in a pediatric cancer and hematologic disease cohort. **A)** The ensemble approach, with automated filtering, identifies significantly fewer fusions compared to individual callers. The number of fusions is plotted as log_10_(x+1) to account for 0 fusions identified in some cases. Callers are sorted by the lowest median number of fusions identified to highest.. **B)** 67 Clinically relevant fusions were identified, represented as a bar graph with decreasing fusions per individual algorithm, highlighting the sensitivity of the ensemble approach compared to individual algorithms. No individual algorithm was able to identify all 67 fusions. **C)** Of the 67 clinically relevant fusions identified, 30 were interchromosomal chimeric (blue), 29 were intrachromosomal chimeric (orange), 3 were loss of function (green), and 5 were promoter swapping (yellow) fusions. **D)** Of the 67 clinically relevant fusions identified, 7 are novel events (red asterisk), while the remaining 60 fusion partners had been described previously in the literature. **E)** A stacked bar graph represents the individual fusion callers that contributed to each clinically relevant fusion.

### Clinical Impact of Fusion Prediction

#### An RBPMS-MET fusion in an infantile fibrosarcoma-like tumor

A female infant presented with a congenital tumor of the right face. Histologically, the tumor consisted of variably cellular fascicles of spindle cells with a nonspecific immunohistochemical staining profile, suspicious for infantile fibrosarcoma. However, the tumor was negative for an *ETV6-NTRK3* fusion, one of the defining features of infantile fibrosarcoma [28]. RNA-Seq of the primary tumor and optimized ensemble pipeline analysis revealed an *RBPMS-MET* fusion as the only consensus call. By contrast, the individual callers identified numerous fusions as follows: Arriba: 16, CICERO: 2142, FusionMap: 29, FusionCatcher: 3907, JAFFA: 1130, MapSplice: 18, and STAR-Fusion: 20 (**Figure 3A, Additional File 3: Table S8**). *RBPMS*, an RNA-binding protein, and *MET*, a proto-oncogene receptor tyrosine kinase, have been identified as fusion partners in a variety of cancers with other genes and as gene fusion partners in a patient with cholangiocarcinoma [29]. Although *MET* fusions are uncommon drivers of sarcoma [30], a *TFG-MET* fusion has been reported in a patient with an infantile spindle cell sarcoma with neural features [31–33]. The interchromosomal in-frame fusion of *RBPMS* (NM_006867, exon 5) to *MET* (NM_000245, exon 15) juxtaposes the RNA recognition motif of RBPMS to the MET tyrosine kinase catalytic domain (**Figure 3B,C**). Given the therapeutic implications of this driver fusion, the fusion was confirmed and reported in the patient’s medical record. The identification of this fusion provided the molecular driver for this tumor, which enabled definitive classification as an infantile fibrosarcoma-like tumor with a *MET* fusion. The patient was initially treated with VAC (vincristine, actinomycin D, and cyclophosphamide) chemotherapy which reduced tumor burden. Surgical resection of the mass was performed with positive margins. Given the presence of a targetable gene fusion, the presence of residual tumor, and the morbidity associated with additional surgery or radiation, the patient was subsequently treated with the MET inhibitor cabozantinib and demonstrated a complete pathological response (**Figure 3D**).

**Figure 3.**
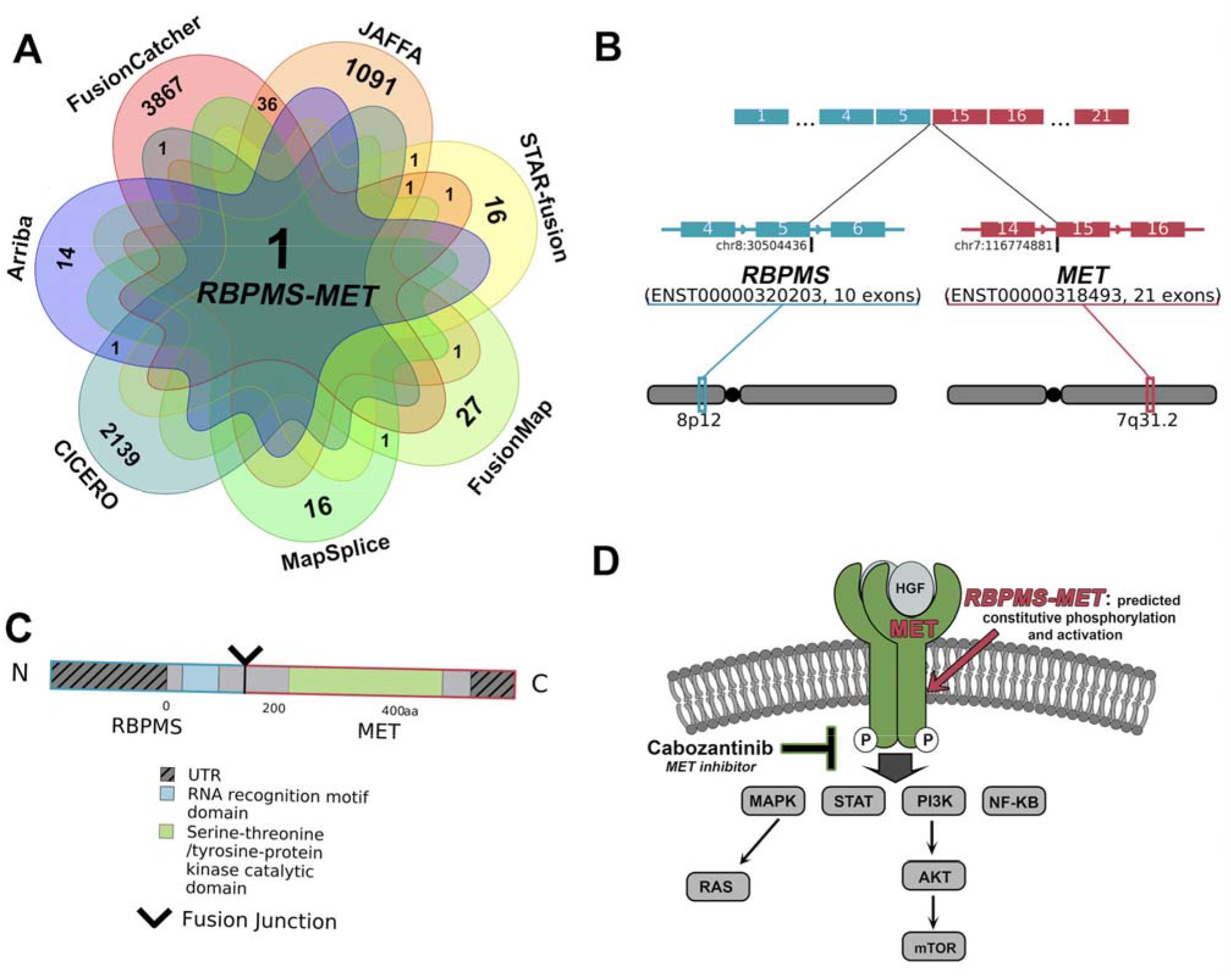
An *RBPMS-MET* fusion identified in a patient with an infantile fibrosarcoma-like tumor. **A)** *RBPMS-MET* fusion was identified by all seven fusion callers in the filtered overlap results. The number of fusions identified by each caller is in the outer VENN diagram sections, while internal numbers indicate overlapping fusions found post-filtering (0 overlaps between callers are not shown). **B)** The *RBPMS-MET* fusion is an interchromosomal event, occurring between 8p12 and 7q31.2 and joining exon 5 of *RBPMS* (blue) to exon 15 of *MET* (red). **C)** The fusion protein product includes the RNA recognition motif domain of RBPMS and the tyrosine kinase catalytic domain of MET. **D)** The *RBPMS-MET* fusion is predicted to cause constitutive phosphorylation and activation of MET, targetable using cabozantinib.

### An NTRK1 fusion in an infiltrating glioma/astrocytoma

A 6-month-old female was diagnosed with an infiltrating glioma/astrocytoma, with a mitotic index of 7 per single high-power field (HPF) and a Ki-67 labeling index averaging nearly 20%, indicative of aggressive disease. RNA-Seq of the primary tumor revealed a *BCAN-NTRK1* fusion, identified by five callers as the only consensus fusion output from the optimized ensemble pipeline (**Figure 4A**). This fusion was clinically confirmed by RT-PCR as an in-frame event, resulting from an intrachromosomal deletion of 225kb at 1q23.1, which juxtaposes *BCAN* (NM_021948, exon 6) to *NTRK1* (NM_002529, exon 8) (**Figure 4B,C**). This fusion results in the loss of the ligand binding domain of NTRK1, while retaining the tyrosine kinase catalytic domain, leading to a predicted activation of downstream targets in a ligand-independent manner [34]. Comparison of the normalized read counts from RNA-Seq data revealed elevated *NTRK1* expression, over 7 standard deviations from the mean, relative to *NTRK1* expression for CNS tumors within the NCH cohort (N=138) (**Figure 4D**). This result indicates the use of first generation TRK inhibitor therapies, with recent regulatory approvals, that have exemplary response rates (75%) and are generally well tolerated by patients [34]. Although the patient has no evidence of disease following gross total resection and treatment with conventional chemotherapy, TRK inhibitors may be clinically indicated in the setting of progressive disease given these findings (**Figure 4E**).

**Figure 4.**
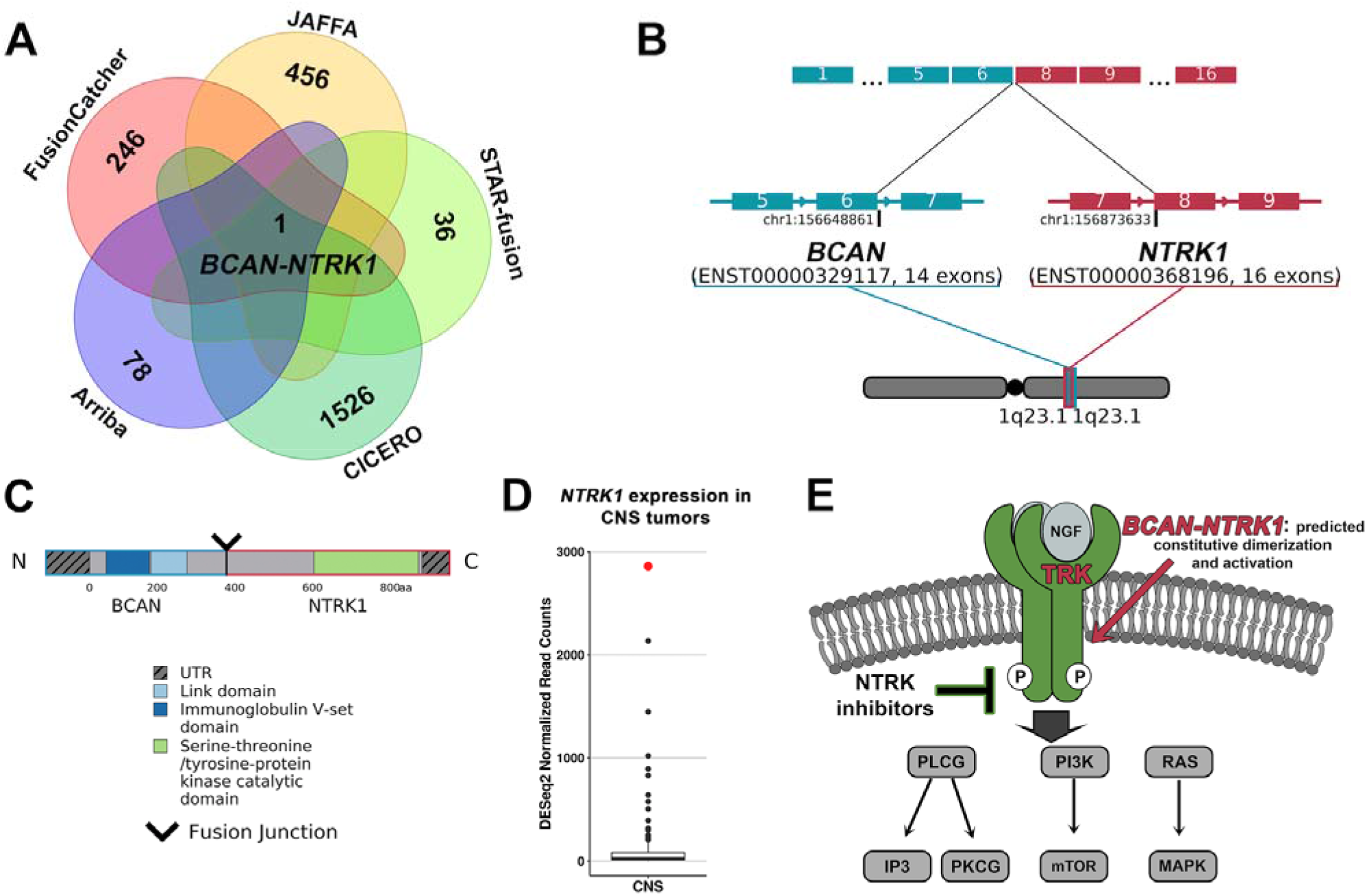
Targetable *NTRK1* fusion identified in an infiltrating glioma. **A)** The *BCAN-NTRK1* fusion was identified by 5 of 7 fusion callers, and was the only fusion returned by the filtered overlap results. Total fusions identified by each caller are shown, FusionMap and MapSplice identified no overlapping fusions that passed filtering (0 overlaps between callers are not shown). **B)** The *BCAN-NTRK1* fusion is an intrachromosomal event occurring on 1q23.1, joining exon 6 of *BCAN* (blue) and exon 8 of *NTRK1* (red). **C)** This fusion results in the juxtaposition of the tyrosine kinase catalytic domain of the *NTRK1* gene to the 5’ end of the *BCAN* gene. **D)** *NTRK1* is highly expressed in this patient (red) compared to CNS tumors (black) in the NCH cohort (CNS tumors: n - 138), with a normalized read count that is 7.70 standard deviations above the mean (131.2). **E)** The *BCAN-NTRK1* fusion is predicted to increase expression and activation of the tyrosine kinase NTRK1, which may be inhibited by TRK inhibitor therapy (green).

### Novel BRAF fusion in a mixed neuronal-glial tumor

A 14-year-old male with a lower brainstem tumor was diagnosed with a low-grade mixed neuronal-glial tumor of unusual morphologic appearance. Tumor histology had features of both ganglioglioma and pilocytic astrocytoma. This tumor was negative for the somatic variant *BRAF* p.V600E, one of the most common somatic alterations associated with gangliogliomas and pilocytic astrocytomas [35], Both the ganglioglioma and pilocytic astrocytoma-like portions of the primary tumor were studied separately by RNA-Seq. A novel *TRIM22-BRAF* fusion was identified in both histologies of the tumor, with fusion overlap results from the ganglioglioma portion represented in **Figure 5A**. *TRIM22-BRAF* was the only consensus fusion output by the optimized fusion detection pipeline, and was clinically confirmed by RT-PCR. *TRIM22* and *BRAF* are novel fusion partners; however, *TRIM22* has been reported with other fusion partners in head/neck squamous cell carcinoma [36]. *BRAF* is a known oncogene that activates the RAS-MAPK signaling pathways, and has been described with numerous fusion partners, including the common *KIAA1549-BRAF* fusion in pediatric low-grade gliomas [35]. This fusion is an interchromosomal translocation occurring between *TRIM22* (NM_006074, exon 2) at 11p15.4 and *BRAF* (NM_004333, exon 9) at 7q34. The resulting protein includes the TRIM22 zinc finger domains and the BRAF tyrosine kinase domain (**Figure 5B,C**). The *TRIM22-BRAF* fusion may lead to constitutive dimerization and activation of BRAF kinase domain, which is indicated by single sample Gene Set Enrichment Analysis (ssGSEA) and is theoretically targetable through RAF, MEK, or mTOR inhibitors (**Figure 5D,E**).

**Figure 5.**
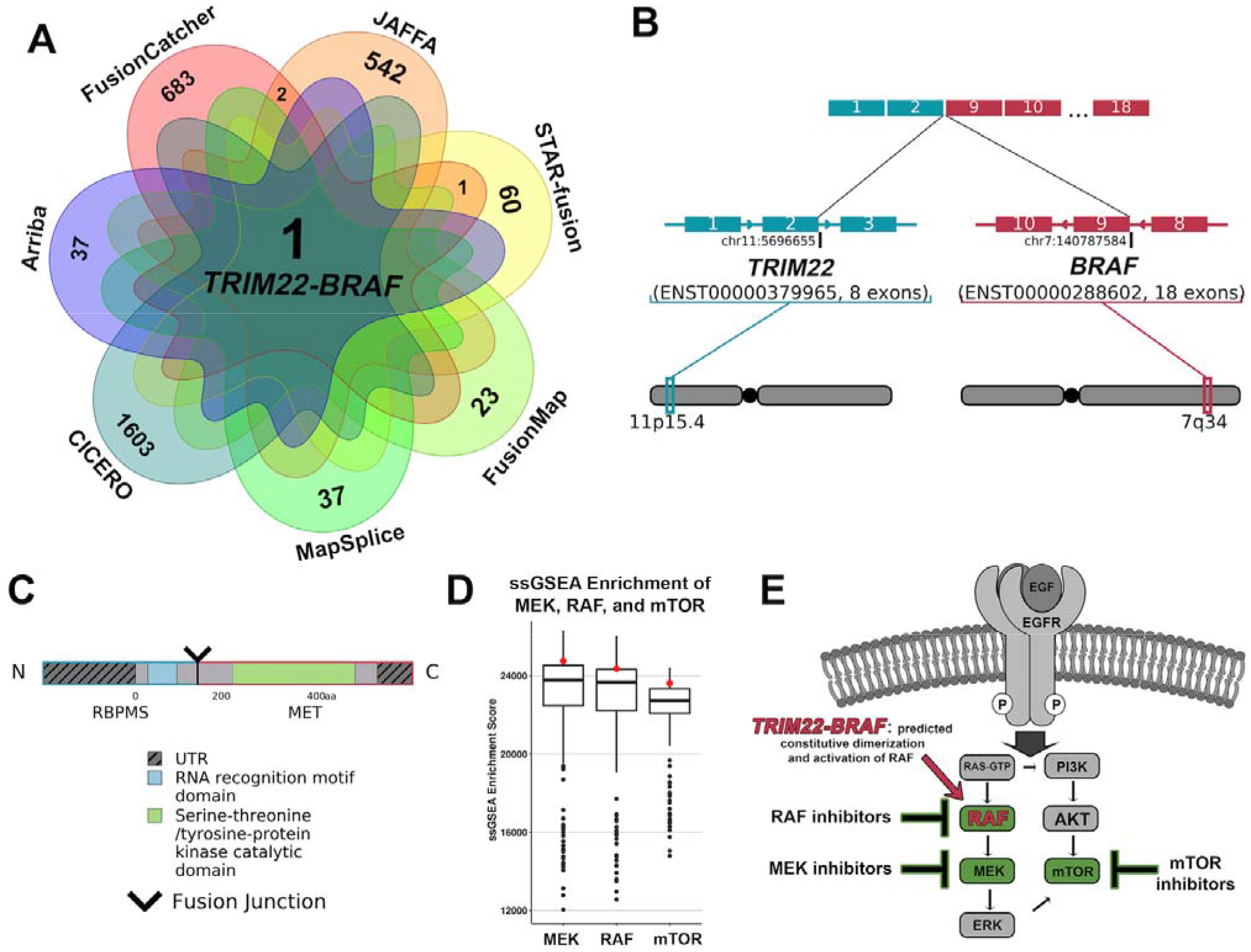
Identification of a novel *BRAF* fusion in a mixed neuronal-glial tumor. **A)** The *TRIM22-BRAF* fusion was identified by all seven fusion callers and in the filtered overlap results, total fusions identified by each caller and overlapping fusions are shown (0 overlaps between callers are not shown). **B)** The *TRIM22-BRAF* fusion is an interchromosomal event between llpl5.4 and 7q34, joining exon 2 of *TRIM22* (blue) to exon 9 of *BRAF* (red). **C)** The resulting fusion product contains the 5’ TRIM22 zinc finger binding domains and BRAF tyrosine kinase catalytic domain. **D)** Single sample gene set enrichment analysis (ssGSEA) indicates a trend toward an enrichment of the MEK (above the 75^th^ percentile, 0.68 standard deviations above the mean of 22756.87), RAF (above the 75^th^ percentile, 0.60 standard deviations above the mean of 22635.74), and mTOR (above the 75^th^ percentile, 0.72 standard deviations above the mean of 22191.50) upregulated gene sets in the *TRIM22-BRAF* sample (red) compared to the pan-cancer NCH cohort (black) (pan-cancer cohort: n = 229).**E)** The *TRIM22-BRAF* fusion is predicted to cause constitutive dimerization and activation of the BRAF kinase domain, shown in D), which could be targeted by RAF, MEK, and mTOR inhibitors (green).

## Discussion

Fusions play a significant role as common oncogenic drivers of pediatric cancers, and their identification may refine diagnosis, inform prognosis, or indicate potential response to molecularly targeted therapies. We have developed an optimized pipeline for fusion detection that harmonizes results from several fusion calling algorithms, filters the output to remove known false positive results, and evaluates the detected fusions compared to a list of known pathogenic fusions. Testing this pipeline on a reference standard indicated that it outperforms single fusion detection algorithms by reducing the number of false positive calls, producing a smaller number of fusions prioritized by the strength of supporting evidence, and suitable for manual inspection. As such, our pipeline greatly simplifies the interpretation process, enabling our multidisciplinary oncology teams to focus on medically relevant findings.

We tested the optimized ensemble pipeline in a prospective study of 229 pediatric cancer and hematologic disease cases and identified 67 fusions. Of these, the fusions from 50 patients were selected for clinical confirmation by an orthogonal method, in our CAP-accredited, CLIA-validated clinical laboratory. All 50 (100% true positive rate) were confirmed to be true fusion events, and were determined to be of clinical relevance by our multidisciplinary care team, providing a diagnostic yield of over 29% across the cohort. (**Additional File 3: Table S8**). Given the high number of putative fusions observed with any single caller, it can be difficult to manually identify a pathogenic fusion amongst a list of tens, if not hundreds, of output fusions. By taking into consideration the frequency in which each fusion occurs in an internal database, as well as the level of evidence based on the number of callers and number of supporting reads by each caller, one can more confidently remove false positives and identify relevant fusions. While our approach does not remove the necessity of manual curation, which is required to determine true clinical relevance of a fusion, it is able to drastically reduce the number of fusions that must be manually assessed, down to ~4 fusions per case, and provides annotations, including a pathogenicity gene partner score, to ease manual interpretation efforts. Our fully automated pipeline aids in prioritization, filtering, and subsequent knowledge-based analysis, providing a more streamlined and less labor-intensive approach to identify fusions, compared to current fusion identification methodologies, drastically reducing the manual workload required to sort through unfiltered or unprioritized results.

The most frequent fusion identified within our pediatric cancer cohort was *KIAA1549-BRAF* (n=12, frequency= 5.2%; **Figure 2B**)[17]. This fusion is characteristically found in pilocytic astrocytomas, which comprise 8.7% of our pediatric cancer cohort (20 out of 229 cases)[37]. We identified five different sets of *KIAA1549-BRAF* breakpoints within our cohort (**Additional File 1: Figure S8A**). The most common fusion patterns represented in the literature are *KIAA1549* exon *16-BRAF* exon 9 (16-9) or *KĨAA1549* exon *15-BRAF* exon 9 (15-9), and these two breakpoints represent 9 of the 12 *KĨAA1519-BRAF* fusions we identified (**Additional File 1: Figure S8B**) [38, 39]. Three additional previously described sets of breakpoints were also identified, *KIAA1549* exon 16-BRAF exon 11 (16-11; n=1), *KIAA1549* exon 15-BRAF exon 11 (15-11; n=1), and *KIAA1549* exon *13-BRAF* exon 9 (13-9; n=1; **Additional File 1: Figure S8**). While the 16-11 and 15-11 breakpoints occur less frequently than 16-9 or 15-9, they have been well described in the literature [38]; whereas only a single case with 13-9 breakpoints was reported as part of a pilocytic astrocytoma cohort study [40]. *KIAA1549-BRAF* fusions often have low levels of expression, a phenomenon that has been described in the literature and is associated with difficulties in its identification through RNA-Seq based methodologies, which lack fusion product amplification [41]. The ability of the ensemble pipeline to identify *KIAA1549-BRAF* fusions, and others that have very low levels of expression, highlights the sensitivity of our approach. Additionally, a supplementary “singleton” file for fusions that are identified by individual algorithms and on the known fusion list is also output by our approach, allowing users the opportunity to manually interpret singleton results. This approach ensures that fusions on the known fusion list are retained, even with minimal evidence by a single caller.

Our approach has also identified other fusions commonly associated with pediatric cancer, including *EWSR1-FLI1* (n=9), *FGFR1-TACC1* (n=3), *PAX3-FOXO1* (n=3), *C11orf95-RELA* (n=2), COL3A1-PLAG1 (n=2), and *NPM1-ALK* (n=2) (**Figure 2B**). In addition to common fusions, our ensemble pipeline also identified seven novel fusions (**Figure 2B**). Five of the seven novel fusions were confirmed by an orthogonal assay in our clinical lab (**Additional File 3: Table S8**). Chimeric fusions, which include both interchromosomal (n=30) and intrachromosomal (n=29) events, were the most common type of fusion identified within the cohort, however, 5 promoter swapping and 3 loss of function fusions were also identified, highlighting the range of fusions this approach is able to detect (**Figure 2D**).

Running seven different fusion callers is computationally complex, as each has its own set of dependencies and environmental requirements. To overcome this, we utilize modern cloud computing technologies. Most notable, our entire pipeline has been built in an AWS serverless environment, removing the requirement for high performance computing (HPC) clusters, while producing highly reproducible results and enabling pipeline sharing. The use of a serverless environment provides flexibility to deploy and scale applications regardless of the application’s size, without needed concern for the underlying infrastructure. We are also leveraging containers to process the data within the serverless environment, as they can be easily utilized by outside institutions with little to no adjustment to their own environments. Another benefit to the current structure of our approach is the ability to assess output from the individual algorithms in real time, as the ensemble pipeline is automatically run after each individual caller completes, allowing for interpretation of at least 3 of the 7 callers within ~3.5 hours, which can be beneficial in situations that necessitate fast turnaround times (**Additional File 1: Figure S5**). Overall, our novel use of serverless technology provides a robust computational solution that is fully automatable and easy to distribute.

There are numerous benefits to the utilization of this optimized pipeline, in that detected fusion events are agnostic to gene partner, allowing identification of common, rare and novel fusions. In addition, the RNA-Seq data set can be utilized for other types of downstream and correlative analyses, including evaluation of gene expression for loci disrupted by the fusion (**Figure 4D**). Utilization of cohort data to assess outlier gene expression can provide valuable insights into pathway disruptions that may occur due to the gene fusion (**Figure 5D**] and may provide information about disease subtyping.

Our ensemble fusion detection pipeline is customizable, allowing users to select how many and which callers to deploy. This may impact potential cost savings, time-to-result, or permit customization that eliminates specific callers that require excessive compute requirements or run times, as suitable in a clinical diagnostic or research setting. Users can also determine the number of consensus calls required to support fusion prediction, which can reduce the number of fusions to assess manually. Callers with a higher percentage of false positives, FusionCatcher and JAFFA, often overlap in their predictions, leading to an increased average number of fusions output by the ensemble pipeline with a consensus of only two callers; a problem diminished by requiring predictions from at least three callers to overlap. In our study, precision was found to be highest in the three-caller consensus version of the ensemble pipeline (**Table 2; Additional File 2: Table S5**). Another benefit to utilizing different algorithms is the ability to assess supplementary output data, in addition to traditional fusion calling. We have made use of this through the inclusion of the internal tandem duplication (ITD) detection which is performed by CICERO. CICERO has identified 7 clinically relevant ITDs within our cohort, 4 of which we have confirmed using orthogonal assays (**Additional File 1: Table S9**).

Future developments to the pipeline could include a weighting system for each caller, based on the precision and sensitivity of the algorithm and on which callers have overlapping predictions, leading to a more sophisticated prioritization strategy. Additional fusion calling algorithms may also be considered and provided as options for users. The known fusion list can also be modified and tailored to include specific gene pairs, or even single genes of interest, providing another layer of customization. Importantly, through the utilization of a proper internal database for frequency filtering purposes, considering age and/or cancer diagnosis, and with the deployment of the appropriate known fusion list, the ensemble approach could be readily implemented in adult cancer fusion detection. Lastly, not all predictors performed equally, and there was a single unresolvable failure of FusionMap to complete. This failure was likely due to the sequencing depth of the sample, however further analysis is required to determine whether parameter modification would permit completion of FusionMap in this case (**Additional File 3: Table S8**). Importantly, our approach was able to circumvent this failure due to the multi-caller nature of the pipeline. Lastly, there are many modalities of RNA-seq analysis that may be harnessed in future developments of the ensemble fusion detection pipeline, which may include an integrative approach exploiting expression-based analysis and ranking. In summary, the ensemble pipeline provides a highly customizable approach to fusion detection that can be applied to numerous settings, with opportunities for future improvements based on additional features and applications.

## Conclusions

The optimized ensemble fusion detection pipeline provides a highly automated and accurate approach to fusion detection, developed to identify high confidence gene fusions from RNA-Seq data produced from pediatric cancer and hematologic disease samples, and could be readily implemented in adult cancer data analysis. The clinical impact of accurately identifying gene fusions in a given patient’s tumor sample is undeniable, not only in terms of refining diagnoses but also in terms of providing prognostic information that shapes treatment decisions. Furthermore, identification of driver fusions may indicate potential response to targeted therapies for cancer patients. The code for the overlap algorithm utilized in this study is publicly available at our GitHub page (https://github.com/nch-igm/nch-igm-ensemble-fusion-detection).

## Methods

### Description of an internal patient cohort

In total, 229 patients were consented and enrolled onto one of three Institutional Review Board (IRB) approved protocols (IRB17-00206, IRB16-00777, IRB18-00786) and studied at the Institute for Genomic Medicine (IGM) at Nationwide Children’s Hospital (NCH) in Columbus, Ohio. Through the utilization of genomic and transcriptomic profiling, these protocols aim to refine diagnosis and prognosis, detect germline cancer predisposition, identify targeted therapeutic options, and/or to determine eligibility for clinical trials in patients with rare, treatment-refractory, relapsed, pediatric cancers or hematologic diseases, or with epilepsy arising in the setting of a low grade central nervous system (CNS) cancer. Our in-house NCH cohort as studied here, consisted of samples from CNS tumors (n=138), hematologic diseases (n=18), and non-CNS solid tumors (n=73), as represented in **Additional File 1: Figure S2**.

### RNA-Seq of patient tissues

RNA was extracted from snap frozen tissue, formalin-fixed paraffin-embedded (FFPE) tissue, peripheral blood, bone marrow, and cerebral spinal fluid utilizing dual RNA and DNA co-extraction methods originally developed by our group for The Cancer Genome Atlas project [42]. White blood cells were isolated from peripheral blood or bone marrow using the lymphocyte separation medium Ficoll-histopauqe. Frozen tissue, white blood cells, or pelleted cells from cerebrospinal fluid were homogenized in Buffer RLT, with beta-Mercaptoethanol to denature RNases, plus Reagent DX and separated on an AllPrep (Qiagen) DNA column to remove DNA for subsequent RNA steps. The eluate was processed for RNA extraction using acid-phenol:chloroform (Sigma) and added to the mirVana miRNA (Applied Biosystems) column, washed, and RNA was eluted using DEPC-treated water (Ambion). DNAse treatment (Zymo) was performed post RNA purification. FFPE tissues were deparaffinized using heptane/methanol (VWR) and lysed with Paraffin Tissue Lysis Buffer and Proteinase K from the HighPure miRNA kit (Roche). The sample was pelleted to remove the DNA, and the supernatant was processed for RNA extraction with the HighPure miRNA column, followed by DNase treatment (Qiagen). RNA quantification was performed with Qubit (Life Sciences).

RNA-Seq libraries were generated using 100 ng to 1 μg of DNase-treated RNA input, either by ribodepletion using the Ribo-Zero Globin kit (Illumina) followed by library construction using the TruSeq Stranded RNA-Seq protocol (Illumina), or by ribodepletion with NEBNext Human/Mouse/Rat rRNA Depletion kit followed by library construction using the NEBNext Ultra II Directional RNA-Seq protocol (New England BioLabs). Illumina 2×151 paired end reads were generated either on the HiSeq 4000 or NovaSeq 6000 sequencing platforms (Illumina). An average of 104 million read pairs were obtained per sample (range 37M to 380M read pairs).

Following data production and post-run processing, FASTQ files were aligned to the GRCh38 human reference (hg38) using STAR aligner (version 2.6.0c)[43]. Feature counts were calculated using HTSeq, and normalized read counts were calculated for all samples using DESeq2 [44, 45]. Single sample Gene Set Enrichment Analysis (ssGSEA), v10.0.3, was performed on DESeq2 normalized read counts using Molecular Signatures Database (MSigDB) Oncogenic Signatures (c6.all.v7.2.symbols.gmt), which included MEK-upregulated genes (MEK_UP.V1_UP), RAF-upregulated genes (RAF_UP.V1_UP), and mTOR-upregulated genes (MTOR_UP.N4.V1_UP) [46].

### RNA-Seq of SeraCare control reference standards

Seraseq Fusion RNA Mix (SeraCare Inc., Milford, MA) was utilized as a control reference standard reagent to test and optimize the ensemble fusion detection pipeline. This product contains 14 synthetic gene fusions *in vitro* transcribed, utilizing the GM24385 cell line RNA as a background. RNA-Seq libraries were prepared utilizing 500 ng input of neat (undiluted) Seraseq Fusion RNA v2, a non-commercially available concentrated product, as input (SeraCare). RNA-Seq libraries were also prepared using 500 ng input of diluted control reference standard (Seraseq Fusion RNA v3 (SeraCare)), which, as a neat reagent is roughly equivalent to a 1:25 dilution of the v2 product, and of total human RNA (GM24385, Coriell) for the fusion-negative controls. Concentrations of individual fusions in the control reference standard were determined by the manufacturer using a custom fluorescent probe set (based on TaqMan probe design) for each fusion and evaluation by droplet digital PCR. Digital PCR-based concentration data (copies/ul) are available in **Additional File 1: Table SI** for the undiluted sample and **Additional File 1: Table S2** for the diluted sample [47].

Dilutions of the Seraseq Fusion RNA v3 reference standard were performed by mixing with control total human RNA (GM24385, Coriell) for final dilutions of 1:25, 1:50, 1:250, 1:500, 1:2500. We also evaluated undiluted Seraseq Fusion RNA v2. For neat and diluted samples, 500ng input RNA was treated using the NEBNext Human/Mouse/Rat rRNA Depletion kit and libraries were prepared following the NEBNext Ultra II Directional RNA-Seq protocol (New England BioLabs). Paired end 2×151 bp reads were produced using the HiSeq 4000 platform (Illumina). An average of 149 million read pairs were obtained per Seraseq sample (range of 86M to 227M read pairs).

### Optimized Fusion Detection Pipeline

Fusions were detected from paired end RNA-Seq FASTQ files utilizing an automated ensemble fusion detection pipeline that employs seven fusion-calling algorithms described in **Table 1**: Arriba (v1.2.0), CICERO (v0.3.0), FusionMap (v mono-2.10.9), FusionCatcher (v0.99.7c), JAFFA (direct v1.09), MapSplice (v2.2.1), and STAR-Fusion (v1.6.0)[25, 48-51]. STAR-Fusion parameters were altered to reduce the stringency setting for the fusion fragments per million reads (FFPM) from 0.05 to 0.02, while default parameters were retained for all other callers. After fusion calling with each independent algorithm, a custom algorithm written in the R programming language, was used to “overlap,” or align and compare, the unordered gene partners identified by individual fusion callers. The utilization of unordered gene partners allows for fusions to be compared, even if different breakpoints were identified by individual algorithms, and to include reciprocal fusions. Fusion partners identified by at least three of the seven callers are retained and prioritized based on the number of contributing algorithms first and then by the number of sequence reads providing evidence for each fusion. The overlap output retains annotations from the individual callers, including breakpoints, distance between breakpoints, donor and acceptor genes, reads of evidence, nucleotide sequence at breakpoint (if available), frequency information from the database, and whether the identified fusion contains “known pathogenic fusion partners”. If discordant breakpoints are identified across callers for a set of fusion partners, the breakpoints with the most evidence, determined by number of supporting reads, are prioritized in the output.

**Table 1.** Performance comparison of individual fusion calling algorithms. Fusion calling algorithms utilized by the ensemble fusion detection pipeline and their contributions to fusion calling in the NCH pediatric cancer and hematologic disease cohort.

| Tool | Version | Aligner | Reference | Average Fusions Called per Case | Sensitivity (Clinically Relevant Fusions Called out of 67) |
| --- | --- | --- | --- | --- | --- |
| Arriba | v1.2.0 | STAR aligner | Haas *et al.*, 2019 Genome Biol | 54 | 86.6% (58) |
| CICERO | v0.3.0 | candidate SV (structural variant) breakpoints and splice junction | Tian *et al.*, 2020 Genome Biol | 1915 | 89.6% (60) |
| FusionMap | v mono-2.10.9 | GSNAP (Genomic Short-read Nucleotide Alignment Program) - 12mer based | Ge *et al.*, 2011 Bioinformatics | 34 | 88.1% (59) |
| FusionCatcher | v0.99.7c | 4 aligners to identify junctions (Bowtie, BLAT, STAR, and Bowtie2) | Nicorici *et al.*, 2014 bioRxiv | 1558 | 89.6% (60) |
| JAFFA | direct v1.09 | BLAT, uses kmers to selects reads that do not map to known transcripts | Davidson *et al.*, 2015 Genome Med | 1141 | 95.5% (64) |
| MapSplice | v2.2.1 | approximate sequence alignment combined with a local search | Wang *et al.*, 2010 Nucleic Acids Res | 37 | 83.6% (56) |
| STAR-fusion | v1.6.0 | STAR aligner | Haas *et al.*, 2019 Genome Biol | 72 | 94.0% (63) |

The fusions are filtered by the following steps (**Figure 1A**). Read-through events, which occur between neighboring genes and are typically identified in both healthy and disease states, are not expected to impact cellular functions [12, 24]. This type of fusion prediction is a source of false positive results, so we have implemented a filter that removes fusions detected between genes fewer than 200,000 bases apart, that occur on the same strand and chromosome. Recurrent fusions with uncertain biological significance have also been identified in normal tissues. To prevent the inclusion of commonly occurring, benign fusions in our output, a PostgreSQL database was used to filter commonly occurring artifactual fusions. This filter removes any expected fusion artifact with greater than a 10% frequency of detection based on our internal cohort. Lastly, to ensure a high level of confidence in the identified fusions, we utilize a minimum threshold for level of evidence, removing fusions that contain fewer than four reads of support from at least one contributing algorithm.

While filtering can remove false positive results and reduces the time needed to review predicted fusions, it also can remove true positive fusions in certain circumstances. To prevent the inadvertent filtering of known fusions, a known fusion list was developed containing 325 pairs of common fusion partners associated with cancer, as identified in COSMIC and TCGA (**Additional File 1: Table S3**)[27, 52]. To increase sensitivity in the identification of known pathogenic fusions, fusion partners that are on the known fusion list are retained as long as at least two callers have identified the fusion. The ensemble pipeline also outputs a supplementary singleton fusion file, containing fusions identified by a single caller that are on the known fusion list, allowing users to examine low evidence fusions that may be of interest.

To prioritize fusions that contain gene partners commonly found in the known fusion list, we developed the “Gene Partner Predicted Pathogenicity Score” based on the frequency of the individual partners in the known fusion list. Of the 325 fusions on the known fusion list, 38 genes are present as a fusion partner ≥ 3 times (**Additional File 1: Table 4, Figure S3**). The most common partners are *BRAF* and *KMT2A*, which are present as fusion partners 28 times each. To aide prediction of novel, or not well described, pathogenic fusions, we developed a score based on known pathogenic gene partners. This score utilizes the frequency of partners present on the known fusion list. The pathogenic frequency score ranges from 10 (most frequent) to 1 (least frequent, but present at least 3 times):

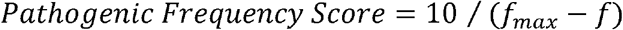

Where *f* is the gene frequency and *f_max_* is the maximum observed frequency. The following annotations are included in the ensemble results if an identified fusion contains one of the 38 common pathogenic gene partners: designation as a known pathogenic gene partner, inclusion of the frequency score (1-10), and gene type based on UniProt description [53].

A knowledge-based interpretation strategy was applied to the filtered list of fusion partners output by the pipeline, including the use of FusionHub [54], to inform clinical relevance, such as diagnostic and/or prognostic information or a potential therapeutic target. Visual assessment of the fusion events was performed by examining RNA-Seq BAM files with Integrated Genome Viewer (IGV). Fusions were also assessed at the DNA level by IGV-based evaluation of gene-specific paired end read alignments from ES or WGS BAM files, for potential evidence of mapping discordance. Clinically relevant fusions were then assayed in our College of American Pathologists (CAP)-accredited clinical laboratory using RT-PCR followed by Sanger sequencing of the resulting products, and/or by Archer FusionPlex Solid Tumor panel (ArcherDx) for clinical confirmation.

### AWS Implementation of the Ensemble Approach

The ensemble fusion detection pipeline is implemented utilizing an Amazon Web Services (AWS) serverless environment (**Additional File 1: Figure S4**). The workflow is initiated via a call to Amazon API Gateway, which passes parameters, including the location of the input FASTQ files, to an AWS Lambda function. The Lambda function initiates the AWS Batch job to load and executes a custom fusion detection Docker image, which launches Arriba, CICERO, FusionMap, FusionCatcher, JAFFA, MapSplice, and STAR-Fusion. We utilize the R5 family of instances for the fusion detection algorithms. Due to the efficiency by which different algorithms are able to multi-thread, each fusion detection tool is allocated 32 virtual CPUs (vCPUs), except for CICERO which is allocated 16 vCPUs and JAFFA which is allocated 8 vCPUs. Using the described allocations, Arriba completes the fastest (~37 minutes) for the runs completed year to date in 2020, followed by FusionMap (~1 hour 12 minutes), STAR-fusion (~3 hours 25 minutes), FusionCatcher (~10 hours 35 minutes), CICERO (~11 hours 49 minutes), MapSplice (~15 hours 2 minutes), and JAFFA (~27 hours 16 minutes), data is summarized in **Additional File 1: Figure S5**. The results from the fusion callers are sent to an AWS S3 output bucket, which invokes AWS Batch to load and execute a Docker image with our overlap script upon completion. This allows for real-time examination of results as each caller finishes, as the overlapping output is updated upon completion of each individual caller, which is particularly advantageous given the long execution times for some of the fusion callers. It is possible to examine results upon completion of the three fastest algorithms within ~3.5 hours, which is of great benefit for cases necessitating fast turnaround times, and complete results are made available by the next day. The overlap Docker image queries and writes to an Aurora PostgreSQL database and performs all necessary filtering. The final results, including annotated filtered and unfiltered fusion lists, are stored in an AWS S3 output bucket for subsequent interpretation. Code for the overlap algorithm is available at our GitHub repository (https://github.com/nch-igm/nch-igm-ensemble-fusion-detection), DOI: 10.5281/zenodo.3950385, and Docker images used to build the pipeline are available upon request.

## Data Analysis and Statistics

Figures were plotted using R version 4.0.2. Statistical analysis was performed by GraphPad Prism 7.0e software. Graphical representation of fusion breakpoints and products were generated using a modified version of INTEGRATE-Vis [55].

## List of abbreviations

AWS: Amazon Web Services
CNS: Central Nervous System
ES: Exome Sequencing
FDR: False Discovery Rate
FFPE: Formalin Fixed, Paraffin Embedded
FFPM: Fusion Fragments Per Million
GSNAP: Genomic Short-read Nucleotide Alignment Program
Heme: Hematologic Diseases
HPF: High Power Field
HPC: High Performance Computing
IGM: Institute for Genomic Medicine
IGV: Integrated Genome Viewer
ITD: Internal Tandem Duplication
NCH: Nationwide Children’s Hospital
QC: Quality Control
RNA-Seq: RNA-Sequencing
ssGSEA: Single Sample Gene Set Enrichment Analysis
vCPU: virtual central processing unit
WGS: Whole Genome sequencing

## Declarations

### Ethics approval and consent to participate

This study was reviewed and approved by the Institutional Review Board (IRB) of The Research Institute at Nationwide Children’s Hospital. Informed consent was obtained from the patients and/or parents for molecular genetic analysis, which included RNA-sequencing. These protocols allowed for return of results from research sequencing studies after confirmation in a CLIA-certified laboratory.

### Availability of data and materials

DNA and RNA sequencing data for this study has been deposited to dbGAP, accession number phs001820.v1.p1

Seraseq fastq files, from the benchmarking studies, have been deposited to the NIH Sequence Read Archive (SRA), accession number PRJNA679580.

Code for the overlap algorithm is available at our GitHub repository (https://github.com/nch-igm/nch-igm-ensemble-fusion-detection), DOI: 10.5281/zenodo.3950385

The Docker image used to run the overlap algorithm is available upon request for running the ensemble pipeline in an AWS serverless environment.

The Docker image used to run previously published fusion detection algorithms is also available upon request for running the ensemble pipeline in an AWS serverless environment.

## Competing interests

### No Competing interests

Stephanie LaHaye, James Fitch, Kyle Voytovich, Adam Herman, Benjamin Kelly, Grant Lammi, Saranga Wijeratne, Kathleen Schieffer, Natalie Bir, Sean McGrath, Anthony Miller, Amy Wetzel, Katherine Miller, Tracy Bedrosian, Kristen Leraas, Ajay Gupta, Bhuvana Setty, Jeffrey Leonard, Jonathan Finlay, Mohamed Abdelbaki, Diana Osorio, Selene Koo, Daniel Koboldt, Vincent Magrini, Catherine Cottrell, Richard Wilson and Peter White.

Elaine Mardis: Qiagen N.V., supervisory board member, honorarium and stock-based compensation.

Daniel Boué: Illumina (ILMN) share holder.

## Authors’ contributions

**SL** analyzed and interpreted fusion data, contributed to development of overlap algorithm and Docker images, contributed to AWS serverless workflow, and wrote the manuscript. **JF** and **KV** contributed development of overlap algorithm and Docker images, contributed to AWS serverless workflow, and contributed to manuscript writing. **AH, BJK, GL,** and **SW** provided data analysis support, designed AWS serverless workflow, and contributed to manuscript writing. **SF** contributed to manuscript revisions and oversaw, organized, and performed data upload to SRA. **KMS** contributed to analysis and interpretation of RNA-Seq results and performed clinical data acquisition. **KM, TAB,** KL and **DK** provided NGS analysis and interpretation for cancer cohort data. **NB, SDM,** and **ARM** perform library preparations and developed laboratory procedures for RNA-Seq processing/QC. **AW** managed RNA-Seq processing and analysis. **KL** managed and coordinated all clinical samples. **DRB, JRL, JLF, MA, DSO, AG, BS,** and **SCK** contributed to the enrollment of patients onto the NCH cancer protocols and provided clinical expertise, **DRB** and **SCK** also contributed pathology materials (fixed or frozen tissues etc.) following QA and/or QC reviews of enrollee pathology. VM oversaw technology development and contributed to the conceptual design of project. **CEC, ERM,** and **RKW** developed, led, and supervised work performed on cancer protocol, contributed to conceptual design of project, contributed to analysis and interpretation of RNA-Seq results, and contributed to manuscript writing and revision. **PW** conceived, designed, and supervised the project, oversaw and contributed to algorithm development, provided support for utilization of AWS and computational resources, and contributed significantly to manuscript writing and revision. All authors read and approved the final manuscript.

## Acknowledgements

We thank the patients and their families for participating in our translational research protocol. We thank the Nationwide Foundation Pediatric Innovation Fund for generously supporting this project.

