## Additional File 1 for "Discovery of Clinically Relevant Fusions in Pediatric Cancer"

-- **ADDITIONAL FILE 1: SUPPLEMENTAL FIGURES AND TABLES** --

**Running title:** Fusion Identification in Pediatric Cancer

FIGURE S1

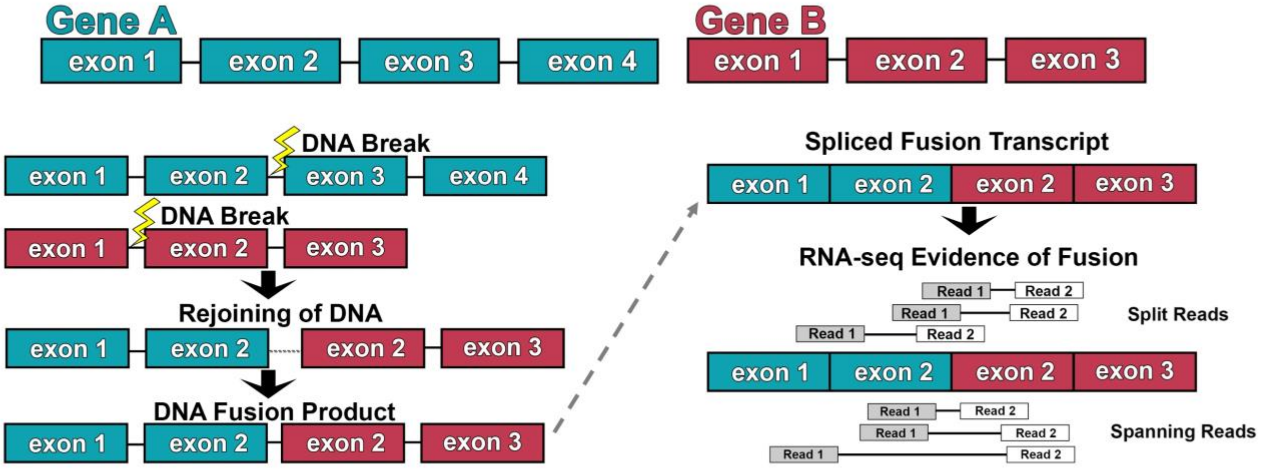


**Figure S1. Fusion occurrence and identification by RNA-Seq.** Fusions occur due to structural changes in genomic DNA, resulting in fusion transcripts. Gene A (blue) and Gene B (red) undergo DNA breaks, rejoin, and create a fusion product. This fusion product undergoes splicing and generates a fusion transcript, which can be identified through RNA-Seq technology. Paired end RNA-Seq identifies fusions through split reads, in which one read includes the junction of the fusion, or through spanning reads, where read 1 (gray) and read 2 (white) map to different genes.

### FIGURE S2

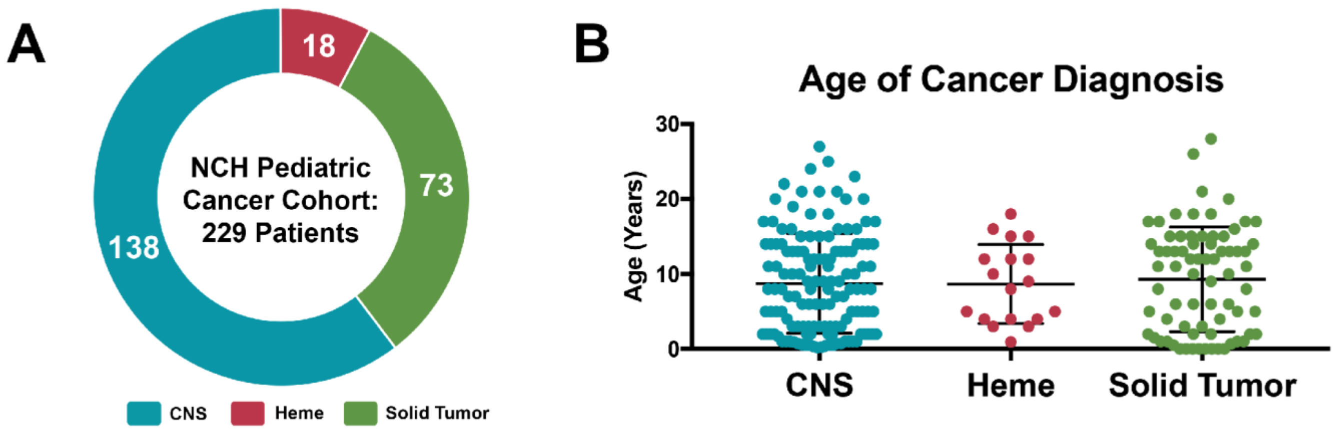


**Figure S2. Pediatric cancer and hematologic disease cohort at Nationwide Children's Hospital. A)** The Nationwide Children's Hospital (NCH) pediatric cancer and hematologic disease cohort, includes three IRB approved studies with a total of 229 patients, consisting of 138 Central Nervous System (CNS) tumors (blue), 18 Hematologic Diseases (Heme; red), and 73 Solid Tumors (green). **B)** The average age of cancer diagnosis for patients included in this study was 8.9 years and is shown broken down by tumor type.

**FIGURE S3**

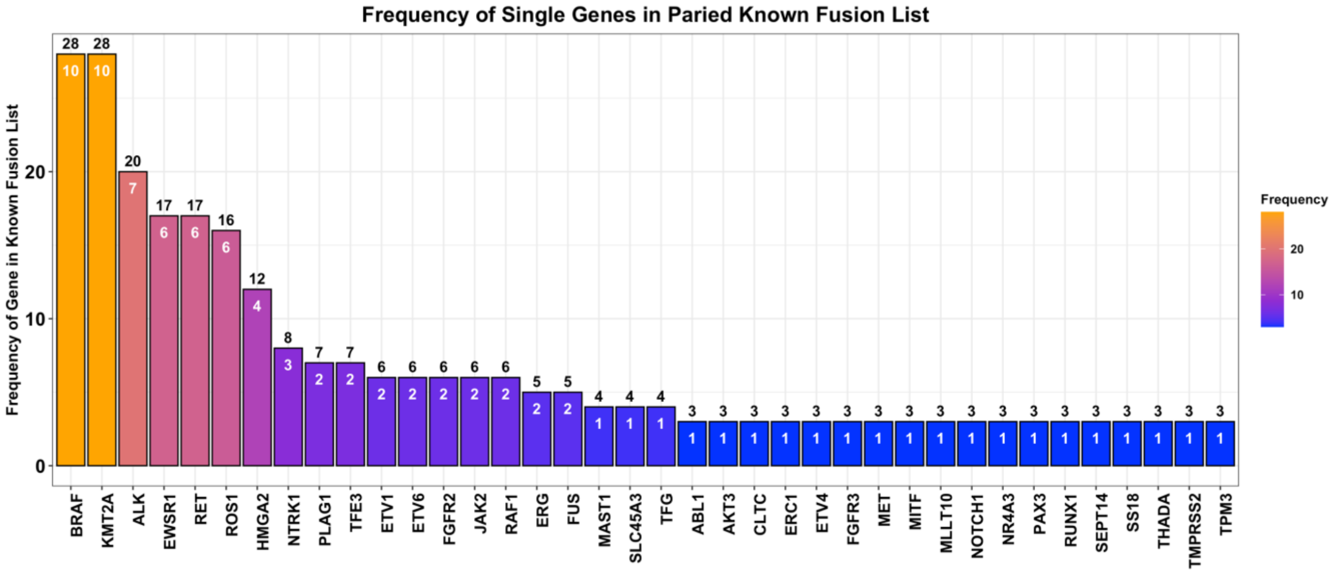


**Figure S3. Frequency and associated pathogenic frequency scores of individual fusion partners in known fusion list.** 38 genes are found commonly (≥3 times) as partners in the known fusion list. Frequency of gene as a partner is shown from high (orange, 28) to low (blue, 3), and associated scores in white are calculated as *Score = 10/*$(f_{max}-f)$, where $f$ is the gene frequency and $f_{max}$ is the maximum frequency observed.

### FIGURE S4

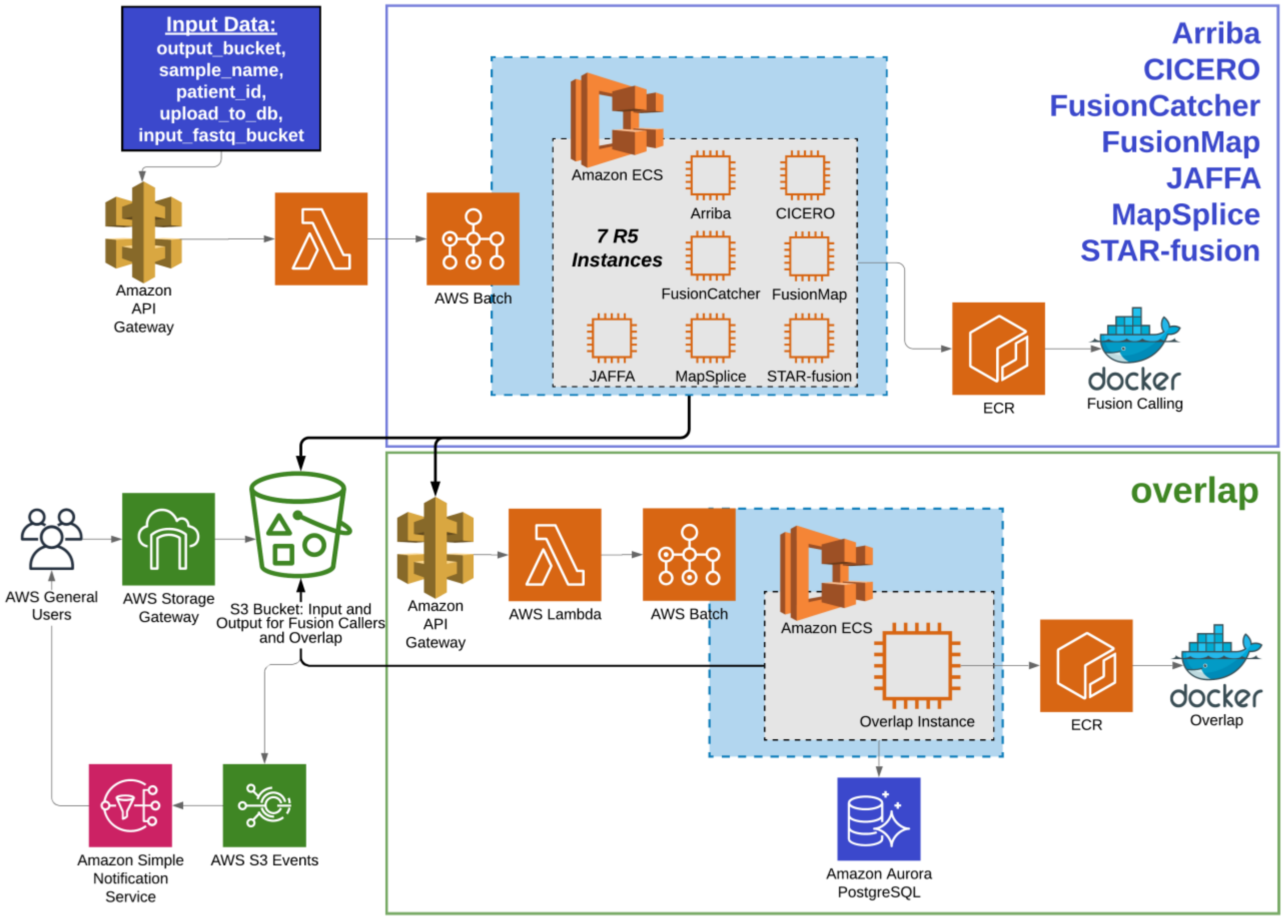


**Figure S4. Ensemble fusion detection architecture diagram for serverless AWS implementation.** Arguments are passed to Amazon API Gateway to invoke Lambda functions, kicking off AWS Batch runs for the Fusion Calling Docker image. Output is sent to an S3 bucket, invoking AWS Batch to kick off the overlap Docker image. The overlap Docker image queries and writes to the Aurora PostgreSQL database and performs all necessary filtering. Final results, including filtered, unfiltered, and singleton results are available in an S3 output bucket.

**Figure S5**

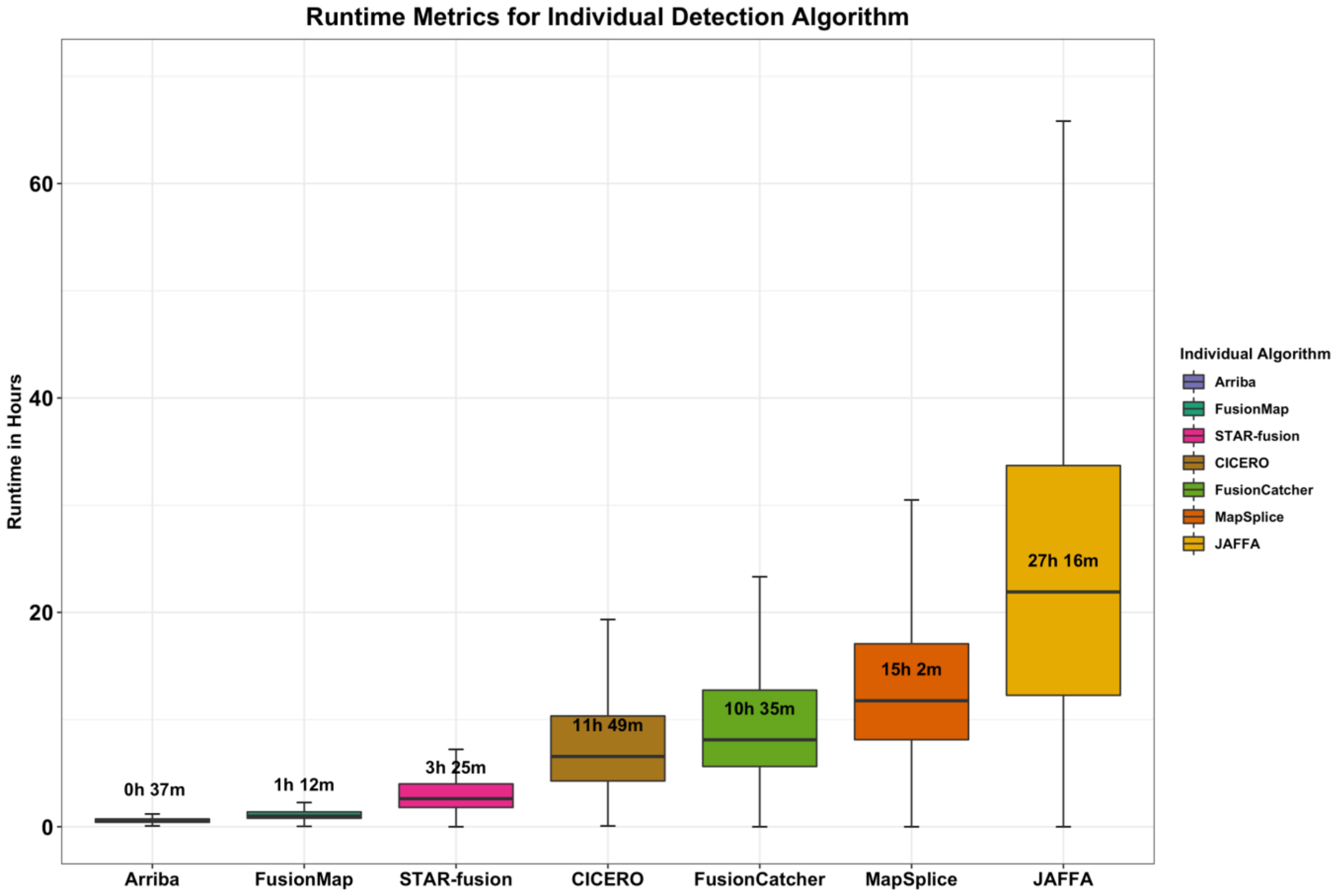


**Figure S5. Averages and distribution of runtimes, in hours, for each fusion detection tool.** Within the tested sample set, Arriba, FusionMap and STAR-Fusion tend to complete in a shorter period of time, even for samples with a large amount of sequencing data, while the remaining 4 tools take substantially longer to complete and have much greater variability in their runtimes. The data shown is from samples processed between January and November 2020, and includes 443 runs of Arriba, 865 runs of FusionMap, 851 runs of STAR-Fusion, 402 runs of CICERO, 827 runs of FusionCatcher, 836 runs of MapSplice and 965 runs of JAFFA. Data is ordered by median runtime, with average runtimes shown.

**FIGURE S6**

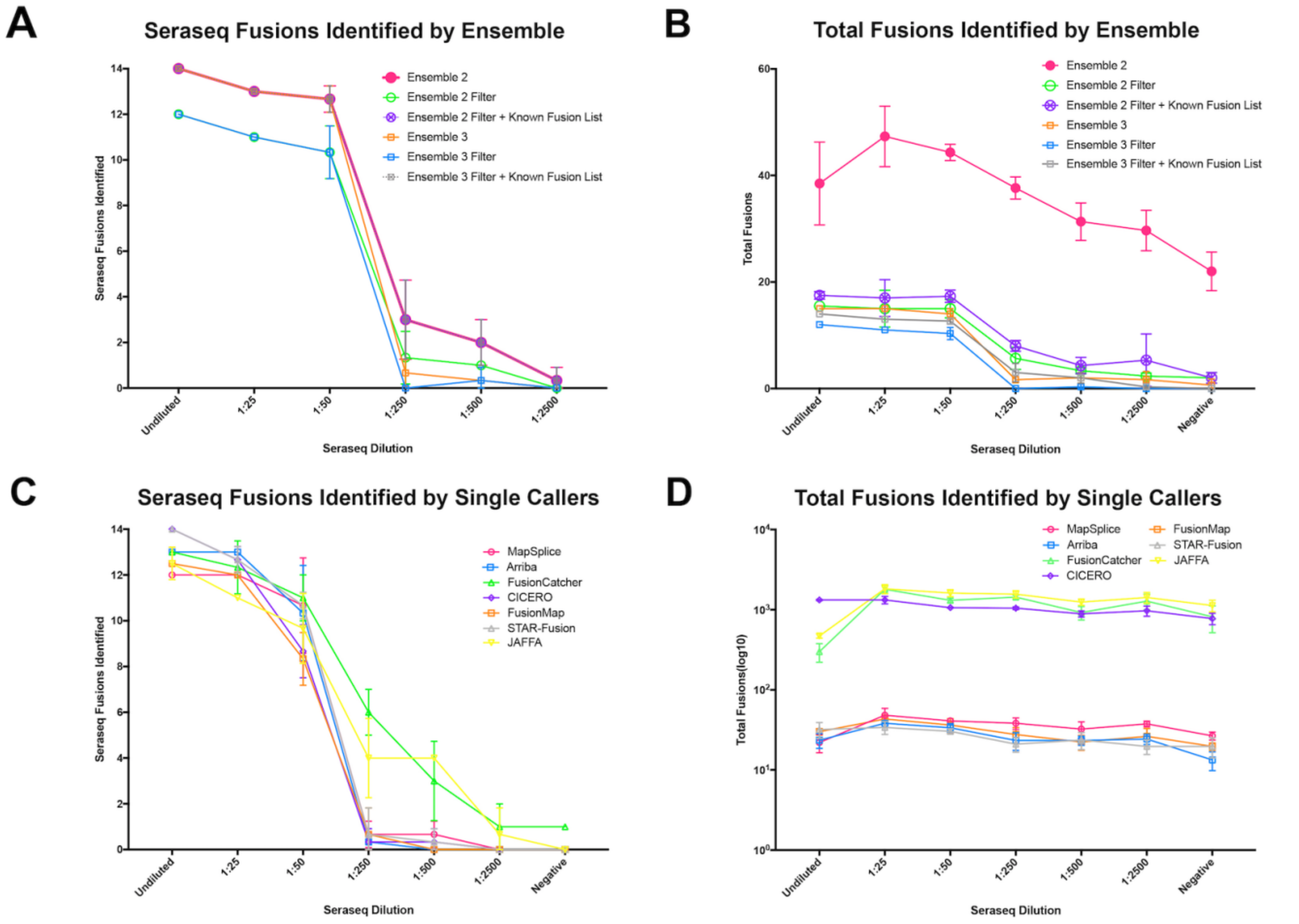


**Figure S6. Limit of detection comparison of ensemble approach and single fusion detection algorithms. A)** Seraseq fusions and **B)** total fusions identified by ensemble approach, including ensemble ≥ 2 callers (Ensemble 2), ensemble ≥ 2 callers with filtering (Ensemble 2 Filter), ensemble ≥ 2 callers with filtering and known fusion list (Ensemble 2 Filter + Known Fusion List), ensemble ≥ 3 callers (Ensemble 3), ensemble ≥ 3 callers with filtering (Ensemble 3 Filter), and ensemble ≥ 3 callers with filtering and known fusion list (Ensemble 3 Filter + Known Fusion List ). **C)** Seraseq fusions and **D)** total fusions identified by each individual caller. Total fusions shown on log10 scale.

**FIGURE S7**

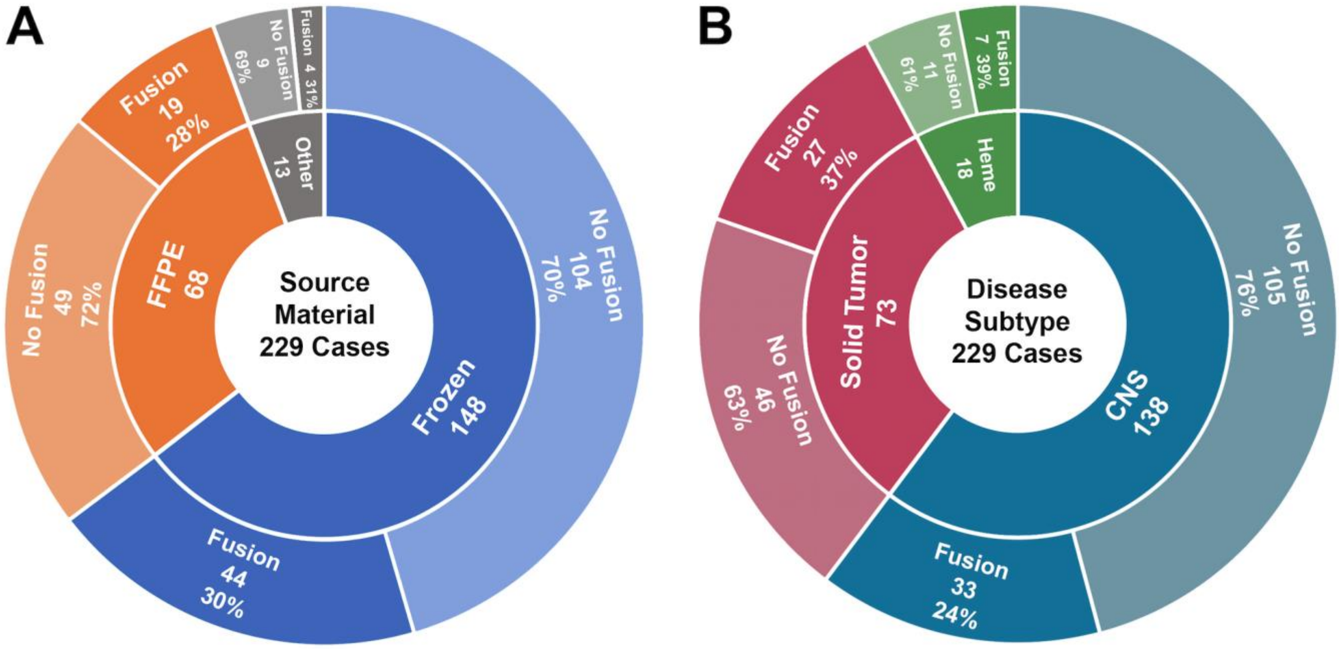


**Figure S7. Fusions identified by source material and disease subtype in 229 pediatric cancer and hematologic malignancies. A)** Independent of sample type, roughly 30% yield was identified across source material types, 44 clinically relevant fusions were identified out of 148 frozen specimens (30% yield), 19 were identified out of 68 FFPE specimens (28% yield), and 4 were identified out of 13 other specimens, which included blood, cerebral spinal fluid, and bone marrow (31% yield). **B)** The cohort is comprised of 138 CNS tumors, 73 solid tumors, and 18 hematologic malignancies. Clinically relevant fusions were identified at the following frequencies by subtype: 33 fusions identified out of 138 CNS specimens (24% yield), 27 fusions identified out of 73 solid tumors (37% yield), and 7 fusions identified out of 18 hematologic malignancies (39% yield). These results highlight the capability of the ensemble approach to identify fusions across pediatric cancer subtypes.

**FIGURE S8**

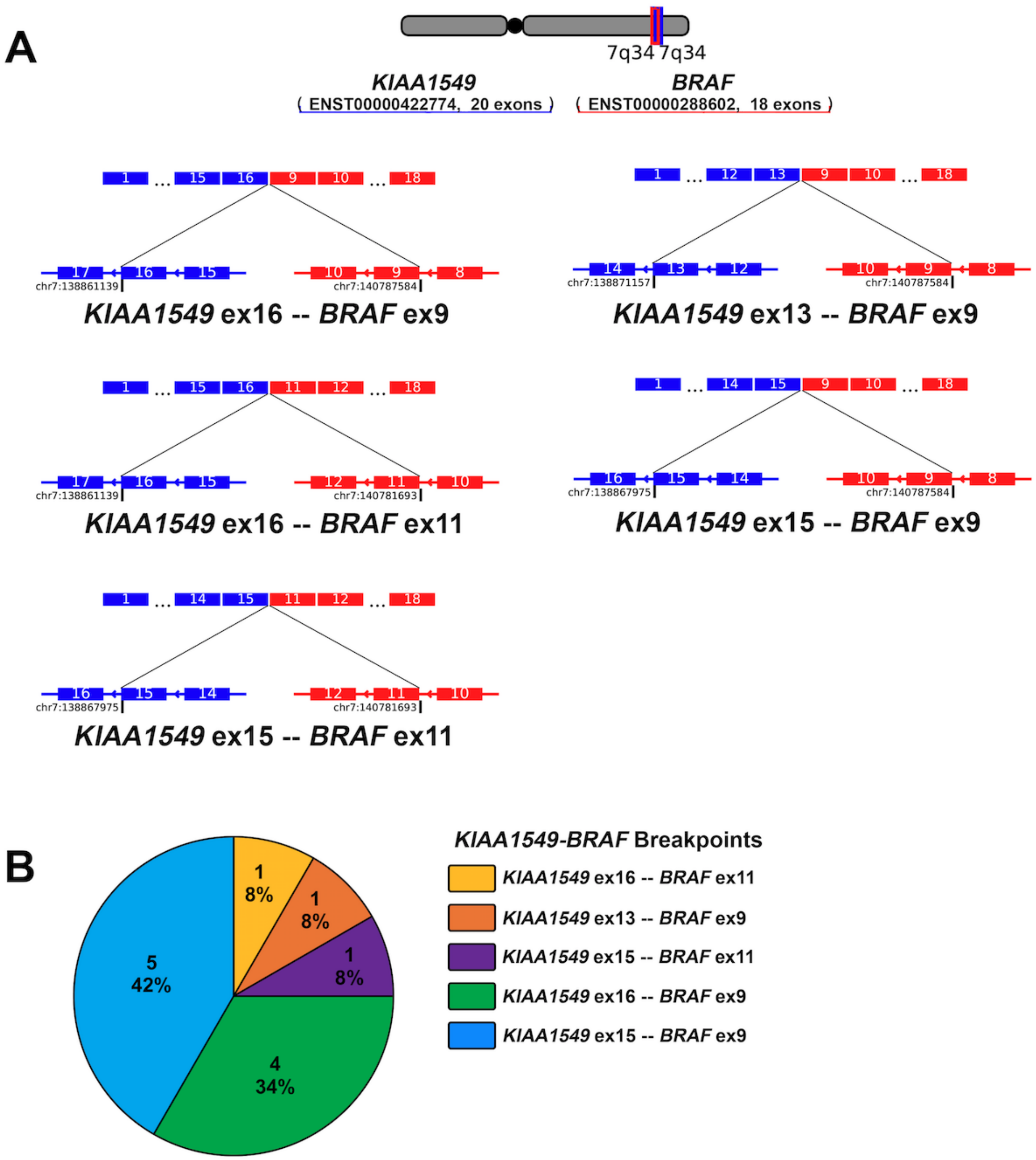


**Figure S8. Breakdown of *KIAA1549-BRAF* fusions in NCH pediatric cancer and hematologic disease cohort. A)** Five different fusion break points were identified in 12 cases, between *KIAA1549* (blue, ENST00000422774/NM_001164665) and *BRAF* (red, ENST00000288602/NM_004333). **B)** Composition of *KIAA1549-BRAF* fusion breakpoints in NCH cancer cohort.

**TABLE S1**

| RNA Fusion | Thermo Fisher Unique Assay ID | Digital PCR copies/μL Member 1 |
|---|---|---|
| *EML4-ALK* | Hs04419883_ft | 18700 |
| *KIF5B-RET* | Hs04396834_ft | 25147 |
| *NCOA4-RET* | Hs04396875_ft | 65813 |
| *CD74-ROS1* | Hs04396897_ft | 21260 |
| *SLC34A-ROS1* | Hs04396942_ft | 35120 |
| *TPM3-NTRK1* | Hs03024839_ft | 29493 |
| *FGFR3-BAIAP2L1* | Hs04396791_ft | 31120 |
| *PAX8-PPARG1* | Hs04396714_ft | 66027 |
| *FGFR3-TACC3* | Hs04421327_ft | 18480 |
| *ETV6-NTRK3* | Hs03024415_ft | 74400 |
| *LMNA-NTRK* | Hs04421343_ft | 32800 |
| *SLC45A3-BRAF* | Hs04396966_ft | 70720 |
| *TMPRSS2-ERG* | AIHSQLA (custom) | 23280 |
| *EGFR-SEPT14* | AJCSWD6(custom) | 50907 |

**Table S1. Digital PCR Results for Undiluted Fusion RNA v2 Dilution Panel RNA.** Results are given as copies of fusion RNA per μL of volume.

**TABLE S2**

| RNA Fusion | Thermo Fisher Unique Assay ID | Digital PCR copies/µL Member 1 |
|---|---|---|
| *EML4-ALK* | Hs04419883_ft | 1347 |
| *KIF5B-RET* | Hs04396834_ft | 539 |
| *NCOA4-RET* | Hs04396875_ft | 797 |
| *CD74-ROS1* | Hs04396897_ft | 891 |
| *SLC34A-ROS1* | Hs04396942_ft | 820 |
| *TPM3-NTRK1* | Hs03024839_ft | 916 |
| *FGFR3-BAIAP2L1* | Hs04396791_ft | 617 |
| *PAX8-PPARG1* | Hs04396714_ft | 1608 |
| *FGFR3-TACC3* | Hs04421327_ft | 1291 |
| *ETV6-NTRK3* | Hs03024415_ft | 879 |
| *LMNA-NTRK* | Hs04421343_ft | 643 |
| *SLC45A3-BRAF* | Hs04396966_ft | 804 |
| *TMPRSS2-ERG* | AIHSQLA (custom) | 605 |
| *EGFR-SEPT14* | AJCSWD6(custom) | 829 |

**Table S2. Digital PCR Results for Seraseq Fusion RNA v3 (before dilution).** Results are given as copies of fusion RNA per µL of volume [31]

**TABLE S3**

| Unordered Fusion | COSMIC | TCGA |
|---|---|---|
| *ABI1+KMT2A* | + | |
| *ABL1+BCR* | + | + |
| *ABL1+ETV6* | + | |
| *ABL1+NUP214* | + | |
| *ACE+KPNB1* | | + |
| *ACSL3+ETV1* | + | |
| *ACTB+GLI1* | + | |
| *ACTN4+KMT2A* | + | |
| *AFDN+KMT2A* | + | |
| *AFF1+KMT2A* | + | |
| *AFF3+KMT2A* | + | |
| *AFF4+KMT2A* | + | |
| *AGGF1+RAF1* | | + |
| *AGK+BRAF* | | + |
| *AKAP13+RET* | | + |
| *AKAP9+BRAF* | + | |
| *AKT3+ARHGAP30* | | + |
| *AKT3+PLD5* | | + |
| *AKT3+ZEB2* | | + |
| *ALK+ATIC* | + | |
| *ALK+CARS* | + | |
| *ALK+CLTC* | + | |
| *ALK+DCTN1* | + | |
| *ALK+DDX6* | | + |
| *ALK+EML4* | + | + |
| *ALK+GTF2IRD1* | | + |
| *ALK+HIP1* | + | |
| *ALK+KIF5B* | + | |
| *ALK+KLC1* | + | |
| *ALK+MALAT1* | | + |
| *ALK+MSN* | + | |
| *ALK+NPM1* | + | |
| *ALK+PPFIBP1* | + | |
| *ALK+RANBP2* | + | |
| *ALK+STRN* | + | + |
| *ALK+TFG* | + | |
| *ALK+TPM3* | + | |
| *ALK+TPM4* | + | |
| *ALK+VCL* | + | |
| *AP3B1+BRAF* | | + |
| *ARHGAP26+CLDN18* | | + |
| *ARHGAP26+KMT2A* | + | |
| *ARHGAP6+CLDN18* | | + |

| Unordered Fusion | COSMIC | TCGA |
|---|---|---|
| *ARID1A+MAST2* | + | |
| *ASPSCR1+TFE3* | + | |
| *ATF1+EWSR1* | + | |
| *ATF1+FUS* | + | |
| *ATG7+BRAF* | | + |
| *ATP1B4+CHST11* | | + |
| *ATP5L+KMT2A* | | + |
| *BAIAP2L1+FGFR3* | + | |
| *BCL2L11+BRAF* | | + |
| *BCOR+ZC3H7B* | + | |
| *BCR+JAK2* | + | |
| *BICC1+FGFR2* | | + |
| *BIRC6+LTBP1* | | + |
| *BRAF+CCNY* | | + |
| *BRAF+CEP89* | + | |
| *BRAF+CLCN6* | | + |
| *BRAF+ERC1* | | + |
| *BRAF+FAM114A2* | | + |
| *BRAF+FAM131B* | + | |
| *BRAF+FCHSD1* | + | |
| *BRAF+GATM* | + | |
| *BRAF+GNAI1* | + | |
| *BRAF+HERPUD1* | + | |
| *BRAF+KIAA1549* | + | |
| *BRAF+LSM14A* | + | |
| *BRAF+MACF1* | | + |
| *BRAF+MKRN1* | + | + |
| *BRAF+RNF130* | + | |
| *BRAF+RUNDC1* | | + |
| *BRAF+SLC45A3* | + | |
| *BRAF+SND1* | + | + |
| *BRAF+SVOPL* | | + |
| *BRAF+TAX1BP1* | | + |
| *BRAF+TRIM24* | + | |
| *BRAF+ZC3HAV1* | | + |
| *BRAF+ZSCAN30* | + | |
| *BRD3+NUTM1* | + | |
| *BRD4+NUTM1* | + | |
| *C11orf1+SIK3* | | + |
| *C2CD2L+ZNF585B* | | + |
| *C8orf34+MET* | | + |
| *CADM2+MITF* | | + |
| *CADM2+TFEB* | | + |

| Unordered Fusion | COSMIC | TCGA |
|---|---|---|
| *CANT1+ETV4* | + | |
| *CAPZA2+MET* | | + |
| *CBFA2T3+GLIS2* | + | |
| *CBFB+MYH11* | | + |
| *CCDC186+FGFR2* | | + |
| *CCDC6+RET* | + | + |
| *CCNB1IP1+HMGA2* | + | |
| *CD74+NRG1* | + | |
| *CD74+ROS1* | + | + |
| *CDH11+USP6* | + | |
| *CDKN2D+WDFY2* | + | |
| *CDX1+IRF2BP2* | + | |
| *CENPP+WNK2* | | + |
| *CEP170B+KMT2A* | + | |
| *CHCHD7+PLAG1* | + | |
| *CIC+DUX4L1* | + | |
| *CIC+FOXO4* | + | |
| *CIPC+NGFR* | | + |
| *CLCN6+RAF1* | + | |
| *CLIP1+ROS1* | + | |
| *CLTC+ROS1* | | + |
| *CLTC+TFE3* | + | |
| *CNTNAP2+GRM8* | | + |
| *COBL+SEPT14* | | + |
| *COL1A1+PDGFB* | + | |
| *COL1A1+USP6* | + | |
| *COL1A2+PLAG1* | + | |
| *COL21A1+TFEB* | | + |
| *COX6C+HMGA2* | + | |
| *CPSF4L+ERBB4* | | + |
| *CREB1+EWSR1* | + | |
| *CREB3L1+FUS* | + | |
| *CREB3L2+FUS* | + | |
| *CREBBP+KAT6A* | | + |
| *CREBBP+KMT2A* | + | |
| *CRTC1+MAML2* | + | |
| *CRTC3+MAML2* | + | |
| *CTNNB1+PLAG1* | + | |
| *CXorf67+MBTD1* | + | |
| *DDIT3+EWSR1* | + | |
| *DDIT3+FUS* | + | |
| *DDX20+TBX15* | | + |
| *DHH+RHEBL1* | + | |
| *DHX33+NLRP1* | | + |
| *DNAJB1+PRKACA* | + | |

| Unordered Fusion | COSMIC | TCGA |
|---|---|---|
| *DPM1+GRID1* | | + |
| *DVL2+TFE3* | | + |
| *EBF1+HMGA2* | + | |
| *EEFSEC+KMT2A* | + | |
| *EGFR+SEC61G* | | + |
| *EGFR+SEPT14* | | + |
| *EIF3E+RSPO2* | + | |
| *ELAVL3+FGFR3* | | + |
| *ELL+KMT2A* | + | + |
| *EP300+KMT2A* | + | |
| *EPHA3+LCLAT1* | | + |
| *EPHB1+MOBKL1B* | | + |
| *EPS15+KMT2A* | + | |
| *ERBB2+MTSS1* | | + |
| *ERBB4+EZR* | + | |
| *ERC1+RET* | + | + |
| *ERC1+ROS1* | + | |
| *ERG+EWSR1* | + | |
| *ERG+FUS* | + | |
| *ERG+NDRG1* | + | |
| *ERG+SLC45A3* | + | + |
| *ERG+TMPRSS2* | + | + |
| *ESRP1+RAF1* | + | |
| *ETV1+EWSR1* | + | |
| *ETV1+HNRNPA2B1* | + | |
| *ETV1+KLK2* | + | |
| *ETV1+SLC45A3* | + | |
| *ETV1+TMPRSS2* | + | + |
| *ETV4+EWSR1* | + | |
| *ETV4+TMPRSS2* | + | + |
| *ETV6+JAK2* | + | |
| *ETV6+MN1* | + | |
| *ETV6+NTRK3* | + | + |
| *ETV6+PDGFRB* | + | |
| *ETV6+RUNX1* | + | |
| *EWSR1+FEV* | + | |
| *EWSR1+FLI1* | + | |
| *EWSR1+MYB* | + | |
| *EWSR1+NFATC2* | + | |
| *EWSR1+NR4A3* | + | |
| *EWSR1+PBX1* | + | |
| *EWSR1+POU5F1* | + | |
| *EWSR1+WT1* | + | |
| *EWSR1+YY1* | + | |
| *EWSR1+ZNF384* | + | |

| Unordered Fusion | COSMIC | TCGA |
|---|---|---|
| *EWSR1+ZNF444* | + | |
| *EXOSC10+MTOR* | | + |
| *EZR+ROS1* | + | + |
| *FGFR1+PLAG1* | + | |
| *FGFR1+TACC1* | + | |
| *FGFR2+FRK* | | + |
| *FGFR2+OFD1* | | + |
| *FGFR2+SHTN1* | | + |
| *FGFR2+VCL* | | + |
| *FGFR3+TACC3* | + | + |
| *FHIT+HMGA2* | + | |
| *FKBP15+RET* | | + |
| *FLT3LG+RPS11* | | + |
| *FLT4+LBH* | | + |
| *FOXO1+PAX3* | + | |
| *FOXO1+PAX7* | + | |
| *FOXO3+KMT2A* | + | |
| *FOXP1+MITF* | | + |
| *FRMD4B+MITF* | | + |
| *GABBR2+NOTCH1* | + | |
| *GAS7+KMT2A* | + | |
| *GOLGA5+RET* | + | |
| *GOPC+ROS1* | + | |
| *GOSR1+ZNF207* | | + |
| *GPBP1L1+MAST2* | + | |
| *H2AFX+WDR18* | | + |
| *HACL1+RAF1* | + | |
| *HAS2+PLAG1* | + | |
| *HEY1+NCOA2* | + | |
| *HLA-A+ROS1* | + | |
| *HMGA2+LHFPL6* | + | |
| *HMGA2+LPP* | + | |
| *HMGA2+NFIB* | + | |
| *HMGA2+PCBP2* | | + |
| *HMGA2+RAD51B* | + | |
| *HMGA2+SENP1* | | + |
| *HMGA2+TSFM* | | + |
| *HMGA2+WIF1* | + | |
| *HNF1B+NOTCH1* | | + |
| *HOOK3+RET* | + | |
| *IGF2BP3+THADA* | | + |
| *INSL3+JAK3* | | + |
| *IRF2BP2+NTRK1* | | + |
| *JAK2+PAX5* | + | |
| *JAK2+PCM1* | + | |

| Unordered Fusion | COSMIC | TCGA |
|---|---|---|
| *JAK2+SEC31A* | + | |
| *JAK2+SSBP2* | + | |
| *JAZF1+PHF1* | + | |
| *JAZF1+SUZ12* | + | |
| *KDM5A+NUP98* | + | |
| *KIAA1797+p16INK4* | | + |
| *KIF5B+RET* | + | |
| *KMT2A+KNL1* | + | |
| *KMT2A+MLLT1* | + | |
| *KMT2A+MLLT10* | + | + |
| *KMT2A+MLLT11* | + | |
| *KMT2A+MLLT3* | + | + |
| *KMT2A+MLLT4* | | + |
| *KMT2A+MLLT6* | + | |
| *KMT2A+SEPT2* | + | |
| *KMT2A+SEPT5* | + | |
| *KMT2A+SEPT6* | + | |
| *KMT2A+SEPT9* | + | |
| *KMT2A+TET1* | + | |
| *KSR1+TENM1* | | + |
| *KTN1+RET* | + | |
| *LANCL2+SEPT14* | | + |
| *LGR5+NUP107* | + | |
| *LIFR+PLAG1* | + | |
| *LMNA+NTRK1* | + | |
| *LRIG3+ROS1* | + | |
| *LRRC37B+NF1* | | + |
| *LTK+UACA* | | + |
| *MAML3+TCF4* | | + |
| *MAML3+UBTF* | | + |
| *MAST1+NFIX* | + | |
| *MAST1+NUP210* | | + |
| *MAST1+TADA2A* | + | |
| *MAST1+ZNF700* | + | |
| *MEAF6+PHF1* | + | |
| *MECOM+RPN1* | | + |
| *MECOM+RUNX1* | | + |
| *MET+TFG* | | + |
| *MLLT10+PICALM* | | + |
| *MLLT10+PPP2R1B* | | + |
| *MYB+NFIB* | + | |
| *MYO5A+ROS1* | + | |
| *NAB2+STAT6* | + | |
| *NACC2+NTRK2* | + | |
| *NBR1+WSB1* | | + |

| Unordered Fusion | COSMIC | TCGA |
|---|---|---|
| *NCOA1+PAX3* | + | |
| *NCOA2+PAX3* | + | |
| *NCOA4+RET* | + | + |
| *NF1+RAB11FIP4* | | + |
| *NONO+TFE3* | + | |
| *NOTCH1+SEC16A* | + | |
| *NR4A3+TAF15* | + | |
| *NR4A3+TFG* | + | |
| *NRG1+SLC3A2* | + | |
| *NSD1+NUP98* | | + |
| *NTRK1+SQSTM1* | | + |
| *NTRK1+SSBP2* | | + |
| *NTRK1+TFG* | + | + |
| *NTRK1+TP53* | + | |
| *NTRK1+TPM3* | + | |
| *NTRK1+TPR* | + | |
| *NTRK2+QKI* | + | |
| *NTRK3+RBPMS* | | + |
| *NUP214+SET* | + | |
| *NUTM2A+YWHAE* | + | |
| *NUTM2B+YWHAE* | + | |
| *OLR1+XIAP* | | + |
| *PAX8+PPARG* | + | + |
| *PBX1+TCF3* | + | |
| *PCM1+RET* | + | |
| *PDCD6+TERT* | | + |
| *PLAG1+TCEA1* | + | |
| *PML+RARA* | + | + |
| *PPFIBP1+ROS1* | + | |
| *PRCC+TFE3* | + | + |
| *PRKAR1A+RET* | + | |
| *PTPRK+RSPO3* | + | |
| *PWWP2A+ROS1* | + | |
| *RAF1+SRGAP3* | + | |
| *RAF1+TRAK1* | | + |
| *RBM10+TFE3* | | + |
| *RELCH+RET* | + | |
| *RET+SPECC1L* | | + |
| *RET+TBL1XR1* | | + |
| *RET+TRIM24* | + | |
| *RET+TRIM27* | + | + |
| *RET+TRIM33* | + | + |
| *RHOT1+TNKS* | | + |
| *ROS1+SDC4* | + | |
| *ROS1+SHTN1* | + | |
| *ROS1+SLC34A2* | + | + |
| *ROS1+TPM3* | + | |
| *ROS1+ZCCHC8* | + | |
| *RPS2P32+THADA* | | + |
| *RUNX1+RUNX1T1* | + | + |
| *SFPQ+TFE3* | + | + |
| *SLC16A14+TRIP12* | | + |
| *SS18+SSX1* | + | + |
| *SS18+SSX2* | + | + |
| *SS18+SSX4* | + | |
| *STIL+TAL1* | + | |
| *TBL1XR1+TP63* | + | |
| *TERT+TRIO* | | + |
| *TG+THADA* | | + |

**Table S3. Inclusion of known pathogenic fusion partners in a known fusion list to prevent over-filtering of known fusions by ensemble pipeline.** The known fusion list includes a total of 325 fusion partners, 191 of which were identified by COSMIC with at least 1 sample tested for the fusion, 104 were identified as known fusions by The Cancer Genome Atlas (TCGA) as reported by Gao et al., 2018, and 30 were identified by both [36, 37] (fusions commonly found in normal tissue have been removed from this list).

**TABLE S4**

| Gene | Description | Frequency | scores |
|---|---|---|---|
| *BRAF* | serine/threonine-protein kinase | 28 | 10 |
| *KMT2A* | methyltransferase | 28 | 10 |
| *ALK* | tyrosine kinase | 20 | 7 |
| *EWSR1* | transcriptional regulator | 17 | 6 |
| *RET* | tyrosine kinase | 17 | 6 |
| *ROS1* | DNA glycosylase/lyase | 16 | 6 |
| *HMGA2* | transcriptional regulator | 12 | 4 |
| *NTRK1* | tyrosine kinase | 8 | 3 |
| *PLAG1* | transcription factor | 7 | 2 |
| *TFE3* | transcription factor | 7 | 2 |
| *ETV1* | transcriptional activator | 6 | 2 |
| *ETV6* | transcriptional repressor | 6 | 2 |
| *FGFR2* | tyrosine kinase | 6 | 2 |
| *JAK2* | tyrosine kinase | 6 | 2 |
| *RAF1* | serine/threonine-protein kinase | 6 | 2 |
| *ERG* | potassium channel | 5 | 2 |
| *FUS* | transcriptional regulator | 5 | 2 |
| *MAST1* | microtubule protein | 4 | 1 |
| *SLC45A3* | solute carrier | 4 | 1 |
| *TFG* | Oncogene | 4 | 1 |
| *ABL1* | Oncogene | 3 | 1 |
| *AKT3* | endoplasmic reticulum transport | 3 | 1 |
| *CLTC* | kinetochore stabilization | 3 | 1 |
| *ERC1* | regulatory subunit of IKK | 3 | 1 |
| *ETV4* | transcriptional activator | 3 | 1 |
| *FGFR3* | transcriptional activator | 3 | 1 |
| *MET* | tyrosine kinase | 3 | 1 |
| *MITF* | transcription factor | 3 | 1 |
| *MLLT10* | transcriptional regulator | 3 | 1 |
| *NOTCH1* | transcriptional regulator | 3 | 1 |
| *NR4A3* | transcriptional activator | 3 | 1 |
| *PAX3* | transcriptional activator | 3 | 1 |
| *RUNX1* | transcriptional regulator | 3 | 1 |
| *SEPT14* | cytoskeletal GTPase | 3 | 1 |
| *SS18* | transcriptional activator | 3 | 1 |
| *THADA* | adenoma associated | 3 | 1 |
| *TMPRSS2* | protease | 3 | 1 |
| *TPM3* | actin filament associated | 3 | 1 |

**Table S4. Commonly occurring fusion partners in known fusion list are utilized as pathogenicity predictor using a score based on gene partner frequency.** 38 genes are present 3 or more times as partners on the known fusion list (see **Additional File 1: Table S3** for known fusion list). Pathogenic frequency scores were developed to predict pathogenicity of a novel, or not well described, fusion based on gene partners known to be common in well described pathogenic fusions. The pathogenic frequency score ranges from 10 (most frequent) to 1 (least frequent, but present at least 3 times): *Score = 10/(28-gene frequency)*. The annotations that are provided in the ensemble output if one of the 38 common pathogenic gene partners is found, are as follows: designation as a known pathogenic gene partner, inclusion of the frequency score (1-10), and a description of the gene type based on UniProt information.

### TABLE S5 – PLEASE SEE ADDITIONAL FILE 2

**Supplemental Materials Table S5. Precision and sensitivity of ensemble approach in Seraseq fusion reagent RNA-seq data.** Row highlighted blue is the average of undiluted (neat) sample duplicates, which is represented in **Figure 1B**. Rows highlighted green are averages per dilution or negative control triplicates (Additional File 2).

TABLE S6

| | Ensemble ≥3 callers | Ensemble ≥2 callers | FusionMap | MapSplice | Arriba | STAR-fusion | FusionCatcher | JAFFA | CICERO |
|---|---|---|---|---|---|---|---|---|---|
| **Minimum** | 0 | 2 | 17 | 18 | 20 | 16 | 244 | 450 | 811 |
| **25% Percentile** | 1.5 | 3.5 | 23.5 | 30 | 20.5 | 19 | 891.5 | 1205 | 991 |
| **Median** | 5 | 9 | 28 | 38 | 27 | 27 | 1350 | 1518 | 1066 |
| **75% Percentile** | 13 | 17.5 | 38.5 | 42 | 36 | 31.5 | 1485.5 | 1659.5 | 1244.5 |
| **Maximum** | 14 | 21 | 49 | 58 | 39 | 39 | 2005 | 2086 | 1488 |
| **Mean** | 7.12 | 10.76 | 31.06 | 37.47 | 28.00 | 26.47 | 1227.41 | 1409.94 | 1089.65 |
| **Std. Deviation** | 5.96 | 6.69 | 8.84 | 9.47 | 7.06 | 7.08 | 468.18 | 425.34 | 184.25 |
| **Std. Error of Mean** | 1.44 | 1.62 | 2.14 | 2.30 | 1.71 | 1.72 | 113.55 | 103.16 | 44.69 |
| **Lower 95% CI of mean** | 4.05 | 7.33 | 26.51 | 32.60 | 24.37 | 22.83 | 986.69 | 1191.25 | 994.92 |
| **Upper 95% CI of mean** | 10.18 | 14.20 | 35.61 | 42.34 | 31.63 | 30.11 | 1468.13 | 1628.63 | 1184.38 |
| **Caller vs. Ensemble ≥3 Callers p value** | - | 1.86E-07 | 3.87E-11 | 2.89E-09 | 8.63E-11 | 1.77E-12 | 1.00E-08 | 3.34E-10 | 3.39E-14 |
| **Caller vs. Ensemble ≥2 Callers p value** | 1.86E-07 | - | 5.77E-10 | 3.21E-08 | 2.3E-09 | 2.95E-10 | 1.03E-08 | 3.43E-10 | 3.50E-14 |

**Table S6. Individual caller statistics for number of total fusions identified in SeraCare Seraseq fusion reagent RNA-seq data.**

Data from ensemble approach with filtering and whitelist shown. Paired student t tests performed on raw values.

**TABLE S7**

| | Ensemble ≥3 callers | Ensemble ≥2 callers | FusionMap | MapSplice | Arriba | STAR-fusion | FusionCatcher | JAFFA | CICERO |
|---|---|---|---|---|---|---|---|---|---|
| **Minimum** | 0 | 0 | 0 | 0 | 0 | 6 | 7 | 2 | 27 |
| **25% Percentile** | 0.5 | 3.5 | 16 | 16 | 12 | 35 | 315 | 464.5 | 1164.5 |
| **Median** | 2 | 8 | 25 | 29 | 30 | 49 | 551 | 768 | 1604 |
| **75% Percentile** | 4 | 19 | 38 | 49 | 73 | 80 | 1082 | 1241 | 2135.5 |
| **Maximum** | 71 | 695 | 695 | 175 | 515 | 1696 | 38726 | 16352 | 8634 |
| **Mean** | 3.93 | 27.07 | 34.53 | 37.43 | 54.20 | 71.62 | 1555.57 | 1146.45 | 1913.34 |
| **Std. Deviation** | 7.74 | 75.29 | 50.66 | 31.37 | 67.96 | 118.98 | 4346.57 | 1541.67 | 1274.81 |
| **Std. Error of Mean** | 0.51 | 4.97 | 3.36 | 2.07 | 4.49 | 7.86 | 287.23 | 101.88 | 84.24 |
| **Lower 95% CI of mean** | 2.93 | 17.27 | 27.92 | 33.35 | 45.35 | 56.12 | 989.61 | 945.71 | 1747.35 |
| **Upper 95% CI of mean** | 4.94 | 36.88 | 41.14 | 41.52 | 63.05 | 87.11 | 2121.54 | 1347.19 | 2079.33 |
| **Caller vs. Ensemble ≥3 Callers p value** | - | 3.17E-06 | 1.99E-17 | 4.68E-43 | 6.71E-24 | 5.89E-16 | 1.65E-07 | 1.28E-23 | 2.52E-60 |
| **Caller vs. Ensemble ≥2 Callers p value** | 3.17E-06 | - | 0.052 (N.S) | 0.044 | 4.1E-05 | 1.71E-10 | 1.70E-07 | 7.66E-24 | 1.51E-61 |

**Table S7. Individual caller statistics for number of total fusions identified in pediatric cancer and hematologic disease cohort data.** Data from ensemble approach with filtering and whitelist shown. Paired Student t tests performed on raw values, N.S. = not significant.

### TABLE S8 – PLEASE SEE ADDITIONAL FILE 3

**Table S8. Fusions identified in pediatric cancer and hematologic disease cohort** (Additional File 3). **.** [1]Clinical confirmation methods include testing performed in a CAP-accredited, CLIA-validated clinical laboratory.

#### TABLE S9

| CaseID | ITD Identified by CICERO | Orthogonal Confirmation |
|---|---|---|
| **IGMCH0053** | EGFR | Yes (Archer FusionPlex panel) |
| **IGMCH0113** | FGFR1 | Yes (Archer FusionPlex panel; RT-PCR followed by Sanger sequencing) |
| **IGMCH0130** | EGFR | Yes (Archer FusionPlex panel) |
| **IGMCH0144** | FGFR1 | Did not confirm by Archer FusionPlex panel |
| **IGMCH0248** | PDGFRA | Did not confirm by Archer FusionPlex panel |
| **IGMCH0252** | FLT3 | Not tested |
| **JL_17015_M17-2670** | FGFR1 | Yes (Archer FusionPlex panel; RT-PCR followed by Sanger sequencing) |

**Table S9. Internal Tandem Duplications (ITD) identified by CICERO in NCH Cohort.** 7 clinically relevant ITDs were identified within the cohort, 4 of which were confirmed using orthogonal assays. These results highlight the benefit of the overlap approach aggregating and utilizing individual pipeline outputs.
